# DHRS7 Integrates NADP^+^/NADPH Redox Sensing with Inflammatory Lipid Signalling via the Oxoeicosanoid Pathway

**DOI:** 10.1101/2025.02.05.636725

**Authors:** Yanan Ma, King Lam Hui, Yohannes A. Ambaw, Tobias C. Walther, Robert V. Farese, Miklos Lengyel, Zaza Gelashvili, Dajun Lu, Philipp Niethammer

## Abstract

During the innate immune response at epithelial wound sites, oxidative stress acts microbicidal and—mechanistically less well understood—as an immune and resilience signal. The reversible sulfhydryl (SH) oxidation of kinases, phosphatases, and transcription factors constitute the perhaps best-known redox signalling paradigm, whereas mechanisms that transduce metabolic redox cues, such as redox cofactor balance, remain little explored.

Here, using mammalian cells, microsomes, and live zebrafish, we identify DHRS7, a short-chain fatty acid dehydrogenase/reductase (SDR), as conserved, 5-hydroxyeicosanoid dehydrogenase (5-HEDH). Under oxidative stress, DHRS7 consumes NADP^+^ to convert arachidonic acid (AA)-derived 5(S)-HETE into the inflammatory lipid 5-KETE, which activates leukocyte chemotaxis via the OXER1 receptor. While Dhrs7 acts as a NADPH-dependent 5-KETE sink in unstressed, healthy tissue, it promotes rapid, 5-KETE dependent leukocytic inflammation in wounded zebrafish skin.

Thus, DHRS7 epitomizes an underappreciated mode of redox signalling—beyond classic SH oxidation—that leverages NADPH metabolism to generate or quench a paracrine lipid signal. Metabolic redox sensors like DHRS7 might be promising therapeutic targets in diseases characterized by disturbed redox balance.

## Introduction

Oxidative stress is a hallmark of inflammatory, cardiovascular, fibrotic, infectious, and neurologic diseases ^1–3^. Although in excess it can cause toxic damage to biomolecules, oxidative stress also serves fundamental innate immune functions. For example, NADPH oxidases of integumental surfaces and white blood cells produce large amounts of reactive oxygen species (ROS) to kill intruding microbes. Oxidative stress also marks potential sites of pathogen entry, such as wounds, to bring in immune reinforcements ^4,5^. Concomitantly, non-lethal oxidative stress confers host resilience against ROS induced bystander damage by upregulating antioxidant circuits or triggering regenerative responses ^6–89^. To serve as diffusible immune and resilience signals, ROS can reversibly modify cysteines of signalling proteins ^10,11^. In addition, oxidative stress may trigger redox-mediated metabolic signals. For instance, at zebrafish wounds, NOX-mediated ROS generation locally alters the NADP^+^/NADPH cofactor balance *in vivo* ^12,13^. Animals and humans lacking epithelial or myeloid NOX activity are prone to infection, for instance, during chronic granulomatous disease, a hereditary NOX2 deficiency ^14–18^. The NOX deficiency phenotypes cannot be solely explained by a lack of microbicidal ROS function and seem to involve aberrant redox signalling. While SH-based redox signalling has been studied in detail, the contribution of “metabolic” redox cues remains largely unexplored.

Zebrafish are a powerful animal model to study redox biology and conserved inflammatory mechanisms to tissue injury by intravital imaging ^19^. Their innate and adaptive immune repertoire and their responses to infection or wounds largely mirror those of mammals ^20,21^. In zebrafish larvae, Ca^2+^ influx at the wound margin activates the epithelial NADPH oxidase Duox. The enzyme produces H_2_O_2_ whilst consuming NADPH to mediate rapid leukocyte recruitment ^4^. Acute neutrophilic wound inflammation via Duox/H_2_O_2_ was shown to involve both reversible Lyn kinase oxidation and production of 5-KETE (5-oxo-eicosatetraenoic acid, alternative name: 5-oxoETE) ^5,22,23^. 5-KETE serves as a potent leukocyte chemoattractant in fish and primates and is linked to allergic inflammation, infection defence, colitis, and possibly cancer ^23–26^. Our companion study reveals 5-KETE as a phylogenetically conserved signal that protects against ROS-induced gut apoptosis, infection, and inflammation, underscoring its importance in inflammatory and adaptive redox signalling (Lengyel et al., 2025). 5-KETE can be formed via non-enzymatic lipid peroxidation or through NADP^+^-dependent 5-HEDH activity ^27^. Research into this pathway was severely stalled for the past 30 years, because 5-HEDH’s genetic identity has remained mysterious and there are no clear OXER1 orthologs in rodents. We therefore sought to identify the enzyme that generates 5-KETE and determine its physiological function in zebrafish.

## Results and Discussion

Earlier work ^27^ has characterized 5-HEDH as a microsomal lipid dehydrogenase/reductase that interconverts 5(S)-HETE and 5-KETE in an NADP^+^/NADPH-dependent manner (**Fig. 1A, S1A**), without identifying its genetic identity. We confirmed reversible 5-HEDH activity in A549 human lung cancer cells using liquid chromatography – mass spectroscopy (LC–MS) to detect product formation after supplying 5(S)-HETE or 5-KETE as substrates (**Fig. 1B**). Consistent with a low cytoplasmic [NADP^+^]/[NADPH] ratio in unstressed cells, there was a pronounced (∼3–4-fold) bias toward 5-KETE consumption. Based on 5-HEDH’s known cofactor dependence, subcellular localization, and evolutionary conservation, we compiled a high-priority 5-HEDH candidate short-list and tested it by short hairpin RNA (shRNA) knockdown in A549 cells. Only knockdown of DHRS7, a short-chain fatty acid dehydrogenase/reductase (SDR) ^28^, significantly reduced 5-KETE below wild type (*wt*) control levels (**Fig. S1B-C**). By contrast, DHRS3 knockdown led to a mild increase of 5-KETE. As the DHRS7 (but not the DHRS3) knockdown effect was consistent with 5-HEDH activity, we tested whether DHRS7 also promoted 5-KETE reduction. This was the case (**Fig. S1D**).

**Figure 1.**
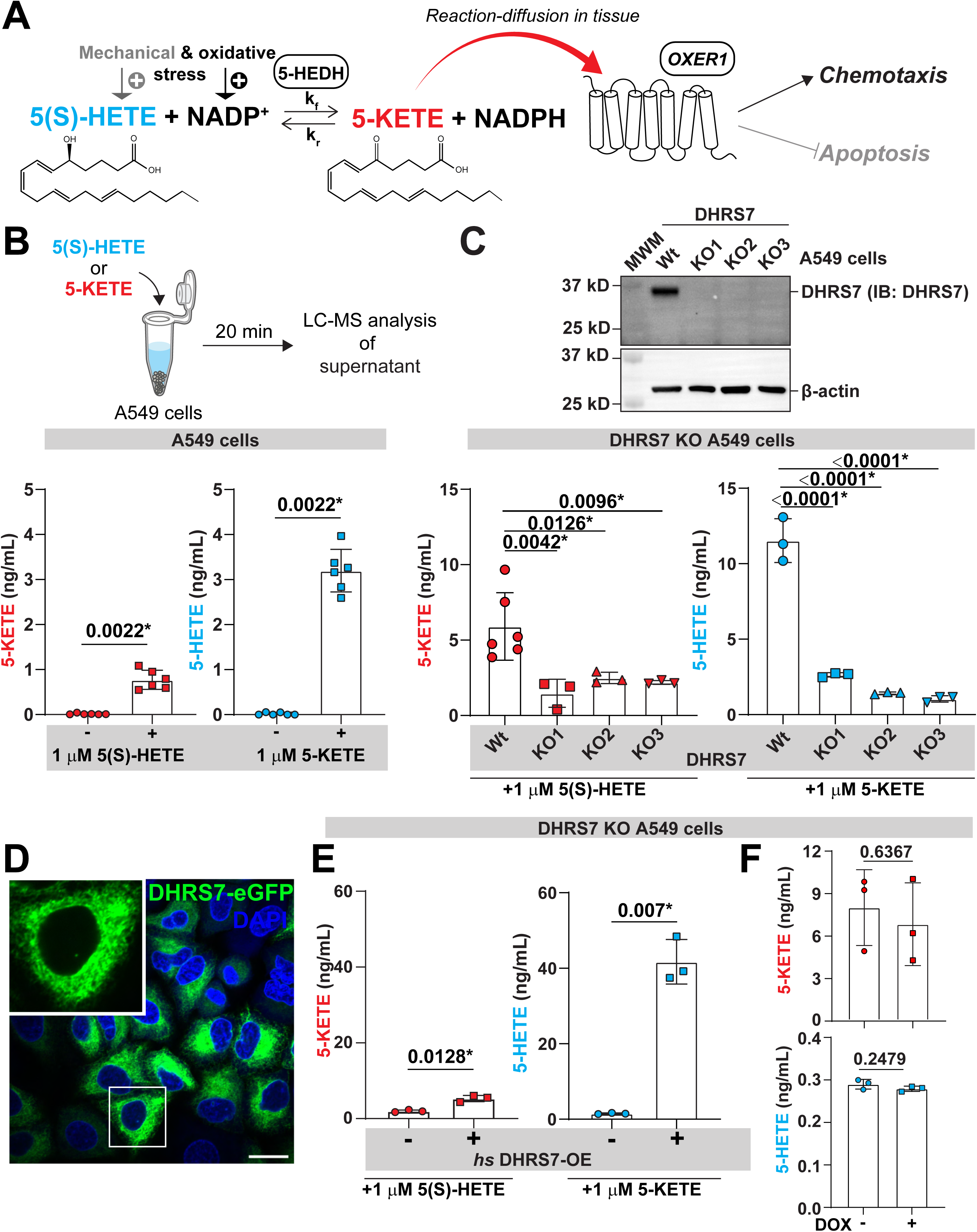
Identification and characterization of 5-HEDH as DHRS7. **(A)** Cartoon scheme of the 5-HEDH reaction and physiological 5-KETE functions. Physical stress leads to the production of 5(S)-HETE, and oxidative stress elevates NADP^+^ levels. The accumulation of these substrates drives the dehydrogenase function of the bidirectional enzyme 5-HEDH, catalysing the conversion of 5(S)-HETE to 5-KETE, generating NADPH. The resulting 5-KETE diffuses into surrounding tissues, exerting chemotactic and anti-apoptotic effects via its GPCR receptor, OXER1. **(B)** Top, a schematic diagram showing the LC-MS assay workflow for measuring 5-HETE or 5-KETE in A549 cells. Bottom, 5-KETE or 5-HETE levels measured by LC-MS upon incubation with 1 μM 5(S)-HETE (left) or 5-KETE (right). P values, unpaired, nonparametric, two tailed Mann-Whitney Test. **(C)** Top, Western blot showing three A549 DHRS7 knockout cell lines. MWM, molecular weight marker. Bottom, LC-MS of A549 DHRS7-KO cells treated with 1 μM 5(S)-HETE (left), or 1 μM 5-KETE (right). P values (left panel), unpaired, parametric, two tailed Welch’s t test. P values (right panel), one-way ANOVA (uncorrected Fisher’s LSD test, two-tailed). **(D)** Subcellular localization of *hs* DHRS7-eGFP in A549 cells. Scale bar, 20 μm. Green, *hs* DHRS7-eGFP. Blue, nuclei (DAPI). Inset, magnified confocal section of indicated cell. **(E)** LC-MS of A549 DHRS7 KO cells ± *hs* DHRS7 overexpression (OE) treated with 1 μM 5(S)-HETE (left), or 1 μM 5-KETE (right). P values, non-paired, parametric, two tailed Welch’s t test. **(F)** 5-KETE or 5-HETE production in DHRS7-KO cells ± 1 μg/mL DOX (without DHRS7 induction). P values, non-paired, parametric, two tailed Welch’s t test. Asterisks, highlight significant changes as compared to respective control (p<0.05).

The DHRS7 nucleotide and protein sequence is conserved from fish to humans (**Fig. S2A, B**). Although experimental DHRS7 structures are yet unavailable, the superposition of AlphaFold predictions for DHRS7 orthologs points to a high structural conservation (**Fig. S2C**). DHRS7 features a NADP(H)-binding Rossman fold ^29^, and localizes to intracellular membranes including those of the endoplasmic reticulum (ER) ^30,31^. Its cofactor preference and localization are consistent with what was reported for 5-HEDH. Whereas DHRS7 was initially proposed to face the ER lumen, subsequent studies demonstrated its cytoplasmic orientation ^32,33,34^ (**Fig. S2D**). DHRS7 is broadly expressed in immune (e.g., macrophages) and non-immune (epithelial, glandular, fibroblast, etc.) cells in various tissues including the prostate, liver, and digestive tract ^31^. Zebrafish single cell RNA sequencing resources suggest that the overall expression pattern is relatively conserved ^35,36^. *In vitro,* DHRS7 possesses reductase activity towards dihydrotestosterone, benzil, and 4,4’-dimethylbenzil ^37^, but its physiological functions and substrates remain unknown. Of note, dehydrogenase activity was not described before for DHRS7.

To independently confirm our shRNA knockdown results, we generated DHRS7 knockout (KO) A549 cell lines using two different single guide RNAs (sgRNAs (**Fig. S2E**). 5-KETE/5-HETE synthesis was significantly reduced in three independent KO clones as compared to *wt* (**Fig. 1C**). Akin to DHRS7’s previously reported endogenous, subcellular localization, doxycycline (DOX)-induced DHRS7 (fused to eGFP, DHRS7-eGFP) localized to endomembranes (**Fig. 1D**) and rescued 5-KETE/5-HETE production (**Fig. 1E**). DOX alone (i.e., without DHRS7) had no such effect (**Fig. 1F**).

To delineate whether or how oxidative stress affects DHRS7-dependent and -independent 5-KETE production (**Fig. 2A-C**), we exposed A549 DHRS7 KO cells ± DOX-induced DHRS7 expression overnight (ON) (**Fig. 2A**) to RSL3, which facilitates lipid peroxidation by inhibiting glutathione peroxidase 4 (GPX4), or imidazole ketone erastin (IKE), which impairs cellular cystine import and glutathione production. Alternatively, cells were briefly subjected to H_2_O_2_, which –unlike RSL3 and IKE—can directly initiate lipid peroxidation in the presence of iron ^38^. H_2_O_2_ also increases the NADP^+^/NADPH ratio (**Fig. S3A**), likely through activating NADP(H) dependent antioxidant circuits that protect cells against irreversible lipid and protein oxidation.

**Figure 2.**
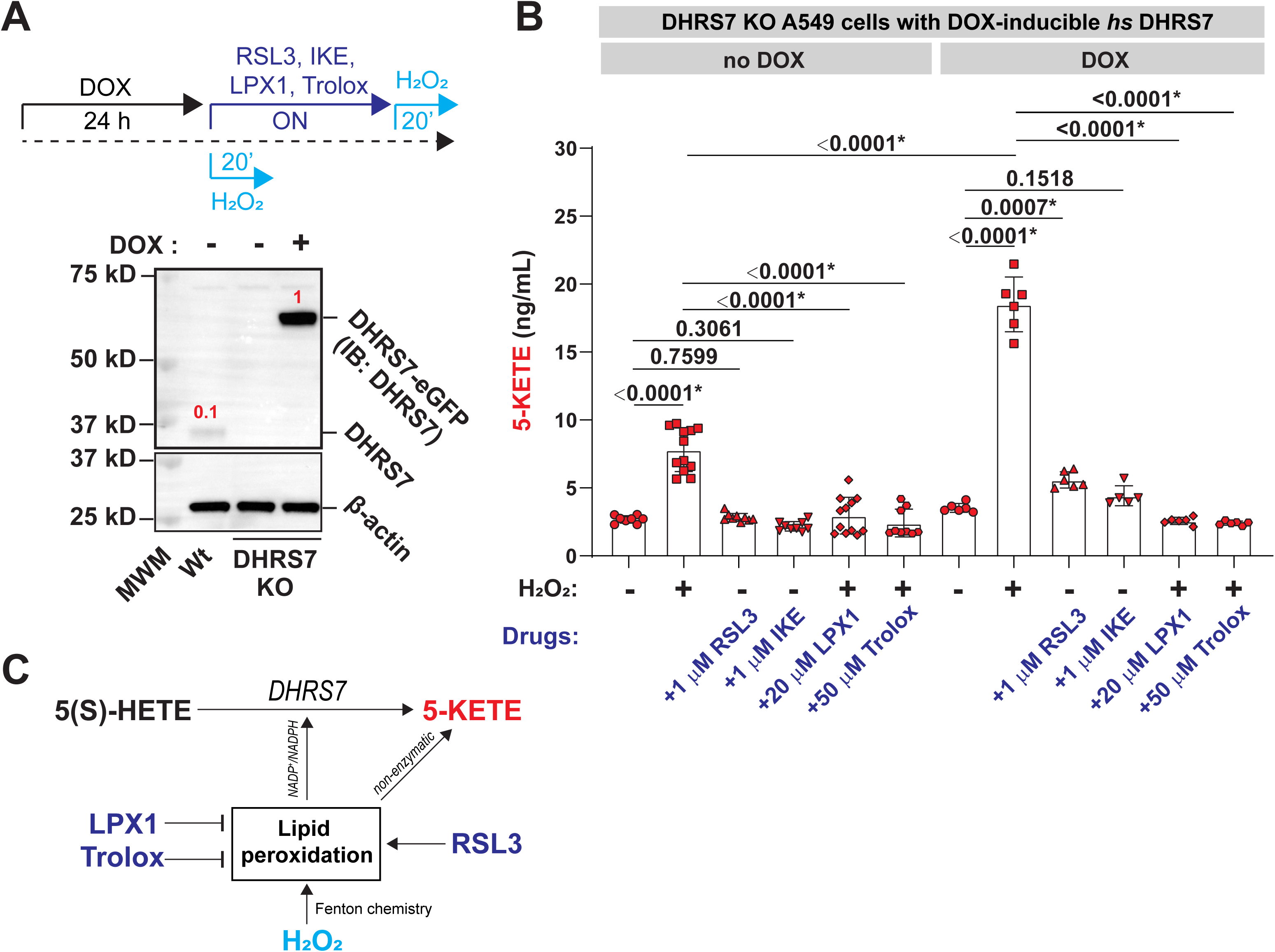
Lipid peroxidation stimulates 5-KETE production through DHRS7. **(A)** Top, experimental scheme. Bottom, western blot showing endogenous DHRS7 and *hs* DHRS7-eGFP expression in *wt* and DHRS7 KO A549 cells upon induction with 0.2 μg/mL doxycycline (DOX). β-actin was used as loading control. Red numbers, relative (β-actin normalized) expression levels as determined by band densitometry. MWM, molecular weight marker. **(B)** LC-MS measurement of 5-KETE at the indicated conditions. P values, one-way ANOVA (uncorrected Fisher’s LSD test, two-tailed). Error bars, SD. **(C)** Scheme of differential drug effects on 5-KETE production. H_2_O_2_ causes oxidative stress, eventually increasing NADP^+^/NADPH through antioxidant circuit activity. As a Fenton reactant, it may also directly promote non-enzymatic 5-KETE production through stimulating free arachidonic acid peroxidation. These processes are counteracted by the ROS scavengers LPX1 (lipophilic) and Trolox (hydrophilic). By contrast, RSL3 (GPX4 inhibitor) inhibits the glutathione dependent termination of the phospholipid peroxidation chain reaction but does not directly stimulate free fatty acid oxidation. Asterisks, highlight significant changes as compared to respective control (p<0.05).

H_2_O_2_ increased 5-KETE levels by ∼ 5 or 17 ng/mL over baseline in DHRS7 KO (**Fig. 2B, left**) or DHRS7 expressing cells (**Fig. 2B, right**), respectively. Thus, despite the presence of non-enzymatic production routes, redox-regulated 5-KETE synthesis by DHRS7 is significant and substantial. Hydrophilic (Trolox) and lipophilic (Liproxstatin-1, LPX1) ROS quenchers completely abrogated H_2_O_2_-dependent enzymatic and non-enzymatic 5-KETE production, presumably through buffering oxidative stress (i.e., limiting NADPH consumption by antioxidant circuits) and scavenging reactive oxygen intermediates involved in non-enzymatic 5-KETE formation. In contrast to H_2_O_2_, 5-KETE induction by RSL3 required DHRS7. We previously showed that lipid peroxidation quenchers (LPX1, Vitamin E) significantly reduce rapid neutrophil recruitment to larval zebrafish wounds *in vivo* ^39^. The above data suggest that this anti-inflammatory effect derives, at least in part, from DHRS7 inhibition. Altogether, these regulatory relationships (**Fig. 2C**) outline a unique role for DHRS7 as metabolic sensor of oxidative stress, non-enzymatic lipid peroxidation, and possibly ferroptosis ^40^.

5-HEDH activity was often studied in microsome suspensions, where redox cofactor concentrations are freely adjustable, and the reaction can be continuously monitored via a NADPH fluorescence readout ^41^. As mentioned above, previous studies suggest that the DHRS7 faces the cytosol ^32,33,34^ (**Fig. S2D**), i.e., the enzyme is exposed to the cytoplasmic cofactor pool. By western blotting and monitoring of NADPH fluorescence in a plate-reader, the endogenous DHRS7 protein and its activity was detected in nuclear, granular (mitochondria, lysosomes), and microsomal fractions of HEK293T cells, but not in the cytosol (**Fig. 3A, S3B-C**). By CRISPR/Cas9, we depleted DHRS7 in HEK293T cells to produce DHRS7 KO microsomes (**Fig. 3B**). Compared to *wt* microsomes, DHRS7 KO microsomes were severely impaired in converting 5(S)-HETE to 5-KETE and vice versa (**Fig. 3C**). However, overexpression of human (*homo sapiens*, *hs*) or zebrafish (*danio rerio*, *dr*) DHRS7 restored both the forward and reverse reaction (**Fig. 3D–F, S3D–E**).

**Figure 3.**
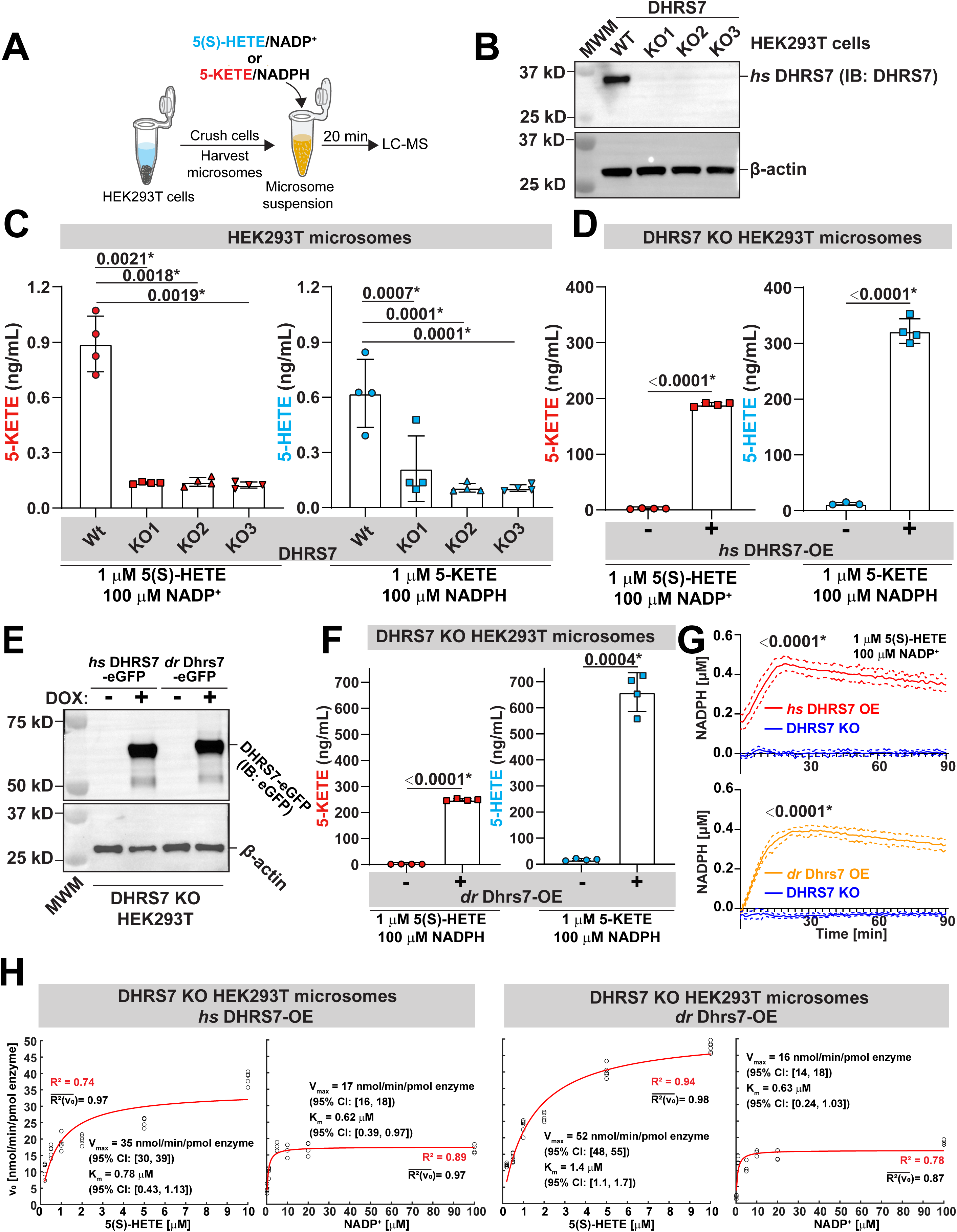
DHRS7’s 5-HEDH activity is phylogenetically conserved. **(A)** Simplified experimental scheme for measuring DHRS7 activity on microsomes by LC-MS. **(B)** Western blot showing three HEK293T DHRS7 KO. MWM, molecular weight marker. **(C)** LC-MS assays with microsomes from HEK293T and *hs* DHRS7-KO cells at the indicated substrate concentrations. P values (left panel), two tailed Welch’s t test. P values (right panel), one-way ANOVA (uncorrected Fisher’s LSD test, two-tailed). **(D)** LC-MS assays with defined amount microsomes/enzyme from *hs* DHRS7-KO (20 μg) and *hs* DHRS7-OE (20 μg, containing 4.39 pmol *hs* DHRS7-eGFP) at the indicated substrate concentrations. P values, unpaired, parametric, two tailed Welch’s t test. **(E)** Western blot confirming equal amounts of *hs* DHRS7-eGFP and drDhrs7-eGFP microsomes in the background of human *hs* DHRS7 knockout cells were used in Figure 2D-2H. **(F)** As in (D) using 4.39 pmol *dr* Dhrs7 instead of 4.39 pmol *hs* DHRS7 (i.e., with 16 μg microsomes containing 4.39 pmol *dr* Dhrs7-eGFP) cells. P values, unpaired, parametric, two tailed Welch’s t test. **(G)** NADPH production assays with microsomes from *hs* DHRS7-KO (10 μg) and *hs* DHRS7-OE (10 μg, containing 2.195 pmol *hs* DHRS7-eGFP) HEK29T cells (top), or from *hs* DHRS7-KO (8 μg) and *dr* Dhrs7-OE (8 μg, containing 2.195 pmol *dr* Dhrs7-eGFP) HEK29T cells (bottom). P values, unpaired, parametric, two tailed, Welch’s t tests comparing the 20 min time points. Solid lines, mean values of data; dotted lines, SD. **(H)** The pseudo-first order Michaelis-Menten reaction parameters were determined by first approximating the initial reaction velocities (v_0_) from the experimentally derived NADPH(5(S)-HETE, t) and NADPH(NADP^+^, t) isotherms by linear fitting their first 20 timepoints. To determine the NADPH(t) isotherms, either one of the substrates of the 5-HEDH dehydrogenase reaction (5(S)-HETE, NADP^+^) was provided in excess while the concentration of the other was altered. [NADPH] was measured using a plate reader assay as a function of time (t). Next, a standard Michaelis Menten model (red line) was fitted to the derived v_0_(5(S)-HETE) *(subpanel-left*) and v_0_(NADP^+^) (*subpanel-right*) isotherms to approximate the pseudo-first order kinetic parameters of the 5-HEDH dehydrogenase reaction. K_m_, Michaelis constant. V_max_, maximum velocity. R^2^(v_0_), average goodness of linear fit for the v_0_ approximation. R^2^, goodness of fit Michaelis-Menten model fit. Brackets, 95% confidence interval (CI). Left subpanel, *hs* DHRS7. Right subpanel, *dr* Dhrs7. Error bars, SD. See methods for further details. Asteriks, highlight significant changes as compared to respective control (p<0.05).

The dehydrogenase reaction of human and zebrafish DHRS7 showed comparable pseudo-first order kinetics when varying [5(S)-HETE] or [NADP^+^] and measuring [NADPH] product formation using the same amount of microsomal *dr* or *hs* DHRS7 (K_m_ ∼0.5–1.5 μM, V_max_ ∼15–50 nmol/min/pmol enzyme; **Fig. 3G–H, S3F**). In contrast, related SDR enzymes (DHRS7B, DHRS7C, DHRS9, and RDH11) showed no significant 5-HEDH activity (**Fig. S4**), suggesting that 5-HEDH activity is a rather specific feature of DHRS7. As expected, NAD^+^ was a less efficient cofactor than NADP^+^ (**Fig. S3F**, middle panel). Besides 5(S)-HETE, DHRS7 also oxidized 5(S)-HEPE, 6-*trans*-12-*epi* LTB4, and 5S,15S-DiHETE, but not 5R-HETE, LTB4, or 12S-HETE (**Fig. S5**). Thus, the allylic 5(S)-hydroxyl group seems to be required and (R)-hydroxyls preclusive for 5-HEDH dehydrogenase activity in line with previous reports on 5-HEDH ^42,43^. Assuming a cytoplasmic NADP(H) pool size of ∼ 3 μM ^12^, its micromolar K_m_ is expected to render DHRS7 an excellent cytoplasmic NADP^+^/NADPH sensor.

However, the physiological role of DHSR7 in animals is still unknown. So, we next examined DHRS7 function in zebrafish using CRISPR/Cas9-mediated knockout (**Fig. S6**). Uninjured mutant tail fins showed elevated 5-KETE and decreased levels of certain prostaglandin-related lipids (e.g., bicyclo-PGE2) compared to *wt* (**Fig. 4A, Table S1**). These data confirm the enzyme’s predicted reductase function in unstressed tissue and suggest some interesting crosstalk with prostaglandin metabolism that warrants further investigation.

**Figure 4.**
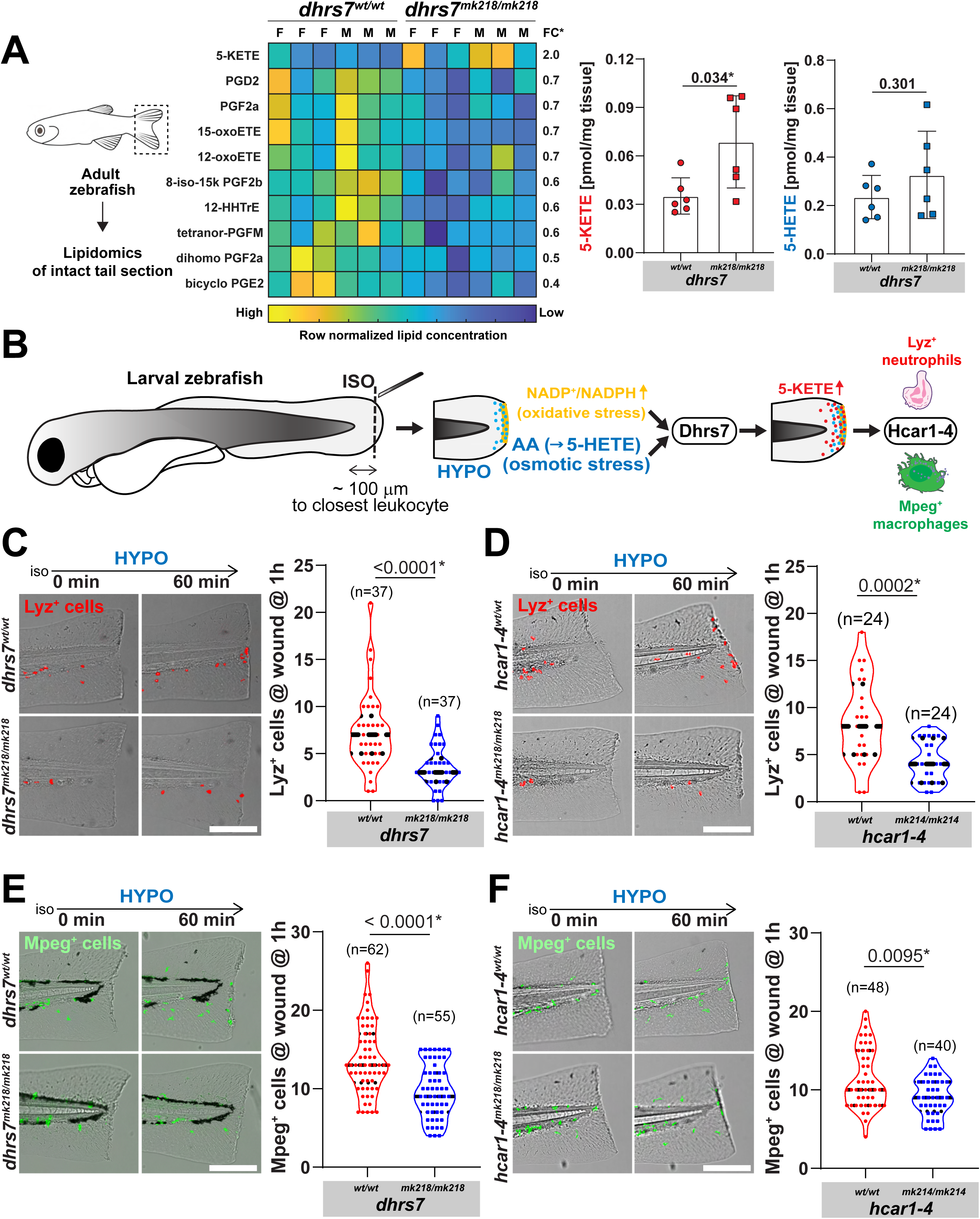
Dhrs7/5-HEDH mediates rapid wound detection in live zebrafish. **(A)** Left, cartoon scheme of lipidomics analysis of adult zebrafish tail fins. Middle, heatmap of all significantly different (*dhrs7^wt^*^/*wt*^ vs. *dhrs7^mk218/mk218^* irrespective of sex, Welch’s t-test or Mann-Whitney U test depending on data normality by Shapiro-Wilk testing) lipid species in the UC San Diego LIPIDMAPS eicosanoid panel, Table S1. FC, fold change. Note, concentrations in the heatmap are row normalized. Right, absolute concentration changes of 5-KETE and 5-HETE between mutant and *wt* animals in the same experiment. P values, parametric, two tailed, Welch’s t-test. Error bars, SD. **(B)** Cartoon depicting the experimental workflow and hypothesis. See main text for further information. ISO, isotonic bathing solution (i.e., blocks 5(S)-HETE generation through inhibiting cPla_2_ activation and AA release; Enyedi et al., 2013, 2016). HYPO, hypotonic, “freshwater-like” bathing solution (i.e., osmotic stress triggers 5(S)-HETE generation through cPla_2_ activation and AA release; Enyedi et al., 2013, 2016). Duox-mediated H_2_O_2_ generation (=oxidative stress) at the wound margin consumes NADPH (Niethammer et al., 2009; Tao et al., 2017; Molinari et al., 2023). Hypothesis: Dhrs7 may integrate the resulting changes in 5(S)-HETE and NADP^+^ to attract neutrophils (*lyz*^+^) and macrophages (*mpeg*^+^) chemotaxis via the OXER1 ortholog Hcar1-4. Macrophage and neutrophil pictograms, NIAID NIH BIOART Source. **(C-D)** Live imaging of neutrophil recruitment to wounds in *wt* siblings and **(C)** *dhrs7^mk218/mk218^* mutant larvae or **(D)** *hcar1-4^mk214/mk214^* mutant larvae (Tg(*lyz*:pm2-mk2) background). Left, representative still images at the indicated times after ISO-HYPO shift. Red, mKate2 expressing neutrophils. Right, quantification. **(E-F)** Live imaging of macrophage recruitment to wounds injury in *wt* siblings and **(E)** *dhrs7^mk218/mk218^* mutant larvae or **(F)** *hcar1-4^mk214/mk214^* mutant larvae (Tg(*mpeg1*:eGFP) background). Left, representative still images at the indicated times after ISO-HYPO shift. Green, eGFP expressing macrophages. Right, quantification. Violin plot error bars, dashed lines denote first quartiles (top line), medians (middle line), third quartiles (bottom line). Parentheses, total number of wounded larvae. P values, unpaired, nonparametric, two tailed Mann-Whitney Test was used in C, D, E, F. Asteriks, highlight significant changes as compared to respective control (p<0.05).

Zebrafish tail-fin wounding simultaneously triggers local, Duox–dependent H_2_O_2_ production that consumes NADPH, and arachidonic acid (AA) release by cPla_2_, providing both the oxidative stimulus and carbon backbone for 5-KETE production (**Fig. S7A**) ^4,12,13,22,44,45^. Of note, cPla_2_ is a nuclear mechanotransducer and its activation, besides Ca^2+^, absolutely requires physical stress, such as, pressure induced nuclear deformation ^22,44,46–48^. We previously showed that cPla_2_ activation is abrogated through isotonic bathing during/after injury and reconstituted by either switching to pure hypotonic solution, or isotonic solution supplemented with the cPla_2_ downstream metabolites AA, 5(S)-HETE or 5-KETE. In the fish, AA and 5(S)-HETE are ultimately converted to 5-KETE by Alox5a ^39^ and 5-HEDH (and/or lipid peroxidation) (**Fig. S7A**). Whereas the redox- and 5-KETE-dependence of wound inflammation is very well established, the mechanisms that integrate oxidative stress (NOX activity) and physical stress (i.e., free fatty acid release) into pro-inflammatory signals remain little understood.

The transient and local dynamics of wound oxidative stress and the large tissue amounts needed for reliable lipid extraction and LC-MS analysis render time-resolved 5-KETE measurements of injured zebrafish tail fins largely infeasible. However, we previously established rapid leukocyte chemotaxis as an excellent *in situ*-proxy for 5-KETE generation at wound sites ^22,23^. So, we counted how many cells expressing fluorescent neutrophil-(*lyz*^+^, red) or macrophage-specific (*mpeg*^+^, green) marker were reaching the wound margin within an hour after switching from isotonic to hypotonic bathing solution to “unfreeze” the wound response (**Fig. 4B**). Orthogonally, we identified generic leukocytes (i.e., neutrophils and macrophages) by transmitted light microscopy irrespective of fluorescent marker expression ^21^. Our homozygous *dhrs7^mk218/mk218^* and *dhrs7^mk219/mk219^* mutants (**Fig. S6B**) showed significantly impaired neutrophil/leukocyte recruitment to wound sites (as compared to *wt* siblings), just like the previously published *hcar1-4*^mk14/mk214^ mutant (**Fig. 4C-D, Video S1-S2, Fig. S7B-D**) ^23^. Interestingly, macrophage recruitment was also inhibited (**Fig. 4E–F, Video S3-S4**). 5(S)-HETE or AA restored wound chemotaxis when the endogenous response was blocked by isotonic bathing conditions, however, only in *wt* not in *dhrs7* mutant animals (**Fig. S7E-F**). Dhrs7 deficient macrophages, but not neutrophils showed impaired wound recruitment towards 5-KETE as compared to *wt* (**Fig. S7G-I**). Per published single cell RNAseq data, Dhrs7 expression strongly clusters with macrophages ^35^. Those phagocytes may harness Dhrs7 as an adaptive chemoattractant sink to steepen shallow 5-KETE gradients “on the go”—an intriguing idea that warrants further testing.

ROS are powerful innate immune toxins that, unlike adaptive immune cells, can unselectively kill known and unknown pathogens (and host cells) alike. Redox signalling may have evolved in part as protective host mechanism to effectively control such dangerous “weapon”. Through enzymes like DHRS7 the host can metabolically (i.e., independent of SH oxidation) couple innate immune cell recruitment to oxidative burst activity of epithelial (Duox) or myeloid (NOX2) NADPH oxidases that indicate/predict pathogenic threats, e.g., at wound sites ^4,12,13^. Besides, redox signalling allows to increase tissue ROS tolerance to selectively protect epithelial barriers, but not the pathogens, against damage. The oxoeicosanoid pathway seems to be essential for both redox signalling arms. As revealed by our companion paper, OXER1 activation, besides being chemotactic ^22,23,27^, counteracts ROS mediated nucleotide oxidation by upregulation of Nudix hydrolases in the gut mucosa (Lengyel et al., 2025).

This highlights 5-KETE as an emerging player in inflammatory disease and cancer of mucosal linings. However, the lack of suitable rodent models (OXER1 ortholog deficiency) has impeded premedical pathway research. By discovering a conserved genetic target—DHRS7—this study now paves the way for new strategies to modulate 5-KETE signalling in human disease. To this end, a structural analysis of DHRS7 would be useful. Although we were, so far, not successful in purifying active enzyme fragments from *E. coli*, more sophisticated approaches are on their way.

Our present data introduce a novel, metabolically driven redox signalling layer—beyond conventional SH oxidation—that may be vital for tissue inflammation and resilience. Future work may unveil whether targeting DHRS7 or related redox-lipid pathways could yield novel therapies for conditions where inflammation and redox imbalance intersect.

## Materials and Methods

### General fish procedures

Adult wild-type and transgenic reporter casper ^49^ zebrafish were maintained at the Memorial Sloan Kettering Cancer Center (MSKCC) Zebrafish Core Facility ^50^, and all experiments were conducted in accordance with institutional animal healthcare guidelines, with approval from the Institutional Animal Care and Use Committee (IACUC) of MSKCC. For wounding assays, larvae aged 2.5 to 3 days post-fertilization (dpf) were anesthetized in isotonic E3 medium (145 mM NaCl, 0.17 mM KCl, 0.33 mM CaCl_2_, 0.33 mM MgSO_4_) containing 0.2 mg/mL ethyl 3-aminobenzoate methanesulfonate (Tricaine/Syncaine, sigma, E10521). At 2.5-3 dpf, the sex of zebrafish larvae cannot be determined, and it is unlikely to influence the biological processes under study. For the adult lipidomics studies, both males and females were used (three biological replicates for each sex and genotype).

### Tail Fin Amputation

For wounding experiments, the tail fin tips of anesthetized 2.5-3 dpf larvae were carefully removed with a surgical microblade (Fine Science Tools, 10318) at the boundary of the notochord, ensuring the notochord remained uninjured ^51^.

### Generation of *dhrs7* mutant zebrafish lines and genotyping

Two independent gRNAs (*dhrs7*-sgRNA-1 and *dhrs7*-sgRNA-2) of zebrafish *dhrs7* (ENSDARG00000003444) were designed and ordered from IDT (integrated DNA Technologies). The Cas9-gRNA ribonucleoprotein complex (single *dhrs7*-sgRNA-1 or a combination of *dhrs7*-sgRNA-1 and *dhrs7*-sgRNA-2) was injected into the cytoplasm of one-cell-stage zebrafish embryos. After the injected F0 larvae matured (2-3 months post-fertilization), individual F0 adults were crossed with wild-type adults to produce F1 progeny. These F1 larvae were then grown to sexual maturity, and genomic DNA was isolated from their tail fins for genotyping. Tail fins were partially amputated, suspended in 250 μL of 50 mM NaOH, and incubated at 95℃ for 10 minutes. Samples were then cooled on ice for 10 minutes, neutralized with 25 μL of 1 M Tris-HCl (pH 8), and vortexed. The genotyping primers are listed in Table S2. PCR products from F1 *dhrs7*-sgRNA-1 adult fish were incubated with FastDigest PvuII (Thermo Fisher Scientific; FD0634) enzyme at 37℃ overnight. The reaction mixture was then separated by agarose gel electrophoresis. F1 heterozygous adults were identified by the presence of three DNA fragments (404, 340, and 68 bp). The 404 bp product represents a mutant allele where the PvuII site has been disrupted by Cas9-induced mutation. This 404 bp band was isolated from the agarose gel and sequenced via Sanger sequencing. F1 heterozygous adult zebrafish with the frameshift mutation of interest (4 bp deletion, *dhrs7*-1, Figure S6) were bred to achieve homozygosity.

PCR products from F1 *dhrs7*-sgRNA-1 and *dhrs7*-sgRNA-2 injected heterozygous adult zebrafish were also separated by agarose gel electrophoresis. The PCR products using the *dhrs7*-sgRNA-1 primers, digested with PvuII, revealed an in-frame mutation with a total 12 bp deletion (8 bp deletion and 4 bp deletion). Meanwhile, the PCR products using the *dhrs7*-sgRNA-2 primers, digested with BseLI (Thermo Fisher Scientific; FD1204), showed a frameshift mutation with a 2 bp deletion. The digested PCR products of *dhrs7*-sgRNA-2 were identified by the presence of three DNA fragments (252, 147, and 107 bp). The 252 bp product represents a mutant allele where the BseLI site has been disrupted by Cas9-induced mutation. This 252 bp band was isolated from the agarose gel and sequenced via Sanger sequencing, confirming the 2 bp deletion. F1 heterozygous adult zebrafish with the frameshift mutation of interest (14 bp deletion (8 bp deletion, 4 bp deletion, 2 bp deletion), *dhrs7-2*, Figure S6) were bred to achieve homozygosity.

### ShRNA

The website http://splashrna.mskcc.org/show_results was used to design shRNA for all the genes mentioned in this manuscript (including HSD17B4 (ENSG00000133835), HSD17B12 (ENSG00000149084), CBR1 (ENSG00000159228), HADH (ENSG00000138796), DHRS3 (ENSG00000162496), PYGB (ENSG00000100994), DHRS1 (ENSG00000157379), DCXR (ENSG00000169738), RDH14 (ENSG00000240857), LMNA (ENSG00000160789), TM7SF2 (ENSG00000149809), RDH13 (ENSG00000275474), PECR (ENSG00000115425), DHRS7 (ENSG00000100612)). The corresponding DNA fragments of these shRNA sequences were ligated into the pSBbi vector using XhoI and EcoRI enzyme restriction sites. Stable cell lines expressing ShRNA were obtained from pSBbi+gene transfected A549 cell lines after selection with blasticidin S hydrochloride (Thermofisher Scientific J61883.FPL).

### Generation of *DHRS7* mutant cell lines

Two independent gRNAs targeting human *DHRS7* (ENSG00000100612), *DHRS7*-sgRNA-1 and *DHRS7*-sgRNA-2 were designed and ordered from IDT (integrated DNA Technologies). The Cas9-gRNA ribonucleoprotein complex, composed of *DHRS7*-sgRNA-1, *DHRS7*-sgRNA-2, Alt-R CRISPR-Cas9 TracrRNA and Alt-R Cas9 enzyme, was introduced into A549 cells or HEK293 cells using the Neon transfection system following the manufacturer’s instructions. Transfected cells were then diluted to single colonies and plated in 96-well plates. After 2-3 weeks of cultivation, the single colonies were expanded in 6-well plates. Western blotting with a human DHRS7 antibody (abcam, ab156021) was performed to confirm DHRS7 knockout in the single colonies. β-actin antibody (sigma, cat#A5441) was used as an internal control. Three independent knockout cell lines were selected for further research.

### NADP^+^/NADPH ratio experiments

HEK293T DHRS7-KO and DHRS7-OE cells were plated in 96-well plates, 1 mM H_2_O_2_ was added to the cells for 20 minutes. Then the NADP^+^/NADPH ratio was detected using the NADP/NADPH-Glo Assay (Promega, G9082). In brief, each well of cells was washed with 50 μl PBS, followed by homogenization with 50 μl of base solution containing 1% DTAB. Next, 50 μl of each sample was transferred to an empty well and mixed with 25 μL of 0.4N HCl. The plate was covered and incubated at 60℃ for 15 minutes, then cooled down to room temperature. To neutralize the acid, 25 μl of 0.5 M Trizma base was added to each well containing acid-treated cells and add 50 μl of HCL/Trizma solution was added to each well containing base-treated samples. Finally, an equal volume of NADP/NADPH-Glo Detection Reagent was added to each well, and the plate was gently shaken and incubated for 30 minutes at room temperature. Luminescence was recorded using Gen5.

### Sample collection and lipid extraction

For the A549 cell culture LC-MS lipids detection, 1,000,000 cells were plated to 6-well plate and incubated at 37℃ overnight. The next day, cells were washed three times by PBS. Then, 1 μM 5-KETE or 1 μM 5(S)-HETE was added to 1.5 ml of Leibovitz’s L15 medium (Thermo fisher Scientific; 21083027) and introduced to each well. The cells were incubated at 37℃ for 20 minutes. The supernatant was collected for the LC-MS analysis.

For the LC-MS lipids detection in *hs* DHRS7-KO A549 cells under different treatments, 500,000 cells were seeded into 6-well plates and incubated at 37℃ overnight. The next day, 1 μg/mL doxycycline was added to induce DHRS7-eGFP expression for 24 hours. On the third day, cells were treated with either 1 μM RSL3, 1 μM IKE, 20 μM LPX1 or 50 μM Trolox overnight (18 hours). After RSL3 or IKE treatment, 1 μM 5(S)-HETE was added for 20 mins. For H_2_O_2_ treatment, 1 mM H_2_O_2_ and 1 μM 5(S)-HETE were added to the medium, with or without 20 μM LPX1 or 50 μM Trolox for 20 min.

Before lipid extraction, 10 μL of a standard solution (0.5 mg/mL 5(S)-HETE-d8 and 0.5 mg/mL 5-KETE-d7) was added to each sample. The eicosanoid extraction was performed with some modifications of an earlier method ^52^. Samples were extracted using Sola Solid Phase Extraction Plates (Thermo Fisher Scientific; A00707). In brief, the columns were conditioned with 3 mL of 100% methanol and then equilibrated with 3 mL of ddH_2_O. Samples were loaded onto the columns, and the sample flow throughs were reloaded to ensure efficient binding. The columns were washed twice with 1 mL of H_2_O: Methanol (90:10 by v/v) to remove impurities. Metabolites/Lipids were eluted with twice 500 μL of 100% methanol. Prior to LC-MS analysis, samples were evaporated using a SpeedVac and reconstituted in 50 μL of H_2_O: acetonitrile-acetic acid (60:40:0.02 v/v/v).

For LC-MS lipids detection in HEK293T microsomes, cells from 10 plates of each genotype were harvested by centrifugation at 3000 × g for 10 min at 4℃. The pellets were resuspended in 50 mL NTE buffer (150 mM NaCl, 15 mM Tris-HCl, 1 mM EDTA, pH 7.5), and centrifuged again at 3000 × g for 10 min at 4℃. The resulting pellets were resuspended in cold cell lysate buffer (0.1 M potassium phosphate buffer pH 7.5, 250 mM sucrose, 50 mM KCl, 1.1 mM EDTA, 0.1 mM DTT, 1:1000 protease inhibitor cocktail (Millipore Sigma, 11873580001)). The cell suspension was lysed using a Dounce homogenizer with 50-60 strokes. To remove unbroken cells, cellular debris, nuclei, and mitochondria, the lysate was centrifuged at 12,000 × g for 20 min at 4℃. Microsomes were isolated by ultracentrifugation at 186,000 × g for 1-2 hours and resuspended in cold resuspension buffer (0.1 M potassium phosphate buffer pH 7.5, 50 mM KCl, 1.1 mM EDTA, 0.1 mM DTT, 20% (v/v) Glycerol, 1:1000 protease inhibitor cocktail). Protein concentration was determined by the Pierce BCA Protein Assay Kits (Thermo Fisher Scientific, PI23227). The concentration of *hs* DHRS7-eGFP and *dr* Dhrs7-eGFP was quantified using eGFP standard.

The enzyme assay was performed as follows: A 1 mL reaction buffer (90 mM potassium phosphate, 40 mM KCl, 0.1 mM DTT, pH 7.5) containing microsomes from *hs* eGFP-KO (20 μg) and *hs* DHRS7-OE (20 μg, containing 4.39 pmol *hs* DHRS7-eGFP), or *hs* DHRS7-KO (16 μg), and *dr* Dhrs7-OE (16 μg, containing 4.39 pmol *dr* Dhrs7-eGFP), 10 μg microsomes (HEK293T, DHRS7-KO1, DHRS7-KO2, DHRS7-KO3), 100 μM NADP^+^ or 100 μM NADPH, 1 μM 5(S)-HETE or 1 μM 5-KETE was incubated at 37℃ for 20 minutes. The reaction was stopped by placing the tubes on ice and adding 10 μL of a reference standard solution (0.5 mg/mL 5(S)-HETE-d8 and 0.5 mg/mL 5-KETE-d7), followed by 4 mL hexane. The aqueous phase was acidified to approximately pH 3.0 by adding 20 μL of 4N HCl to facilitate the extraction of carboxylic acids. Samples were vortexed and centrifuged at 3000 × g for 10 minutes at 4℃ to ensure phase separation and remove precipitated materials. The organic phases were collected, evaporated using a SpeedVac, and reconstituted in 150 μL buffer A (water/acetonitrile/acetic acid, 60/40/0.02, v/v/v).

Whereas absolute 5-KETE/5-HETE baseline levels often varied between individual lipid extractions (e.g., **Fig. 1B** vs. **C**), within the same extraction/LC-MS session, the relative differences were consistent (e.g., **Fig. 1B**: “-” vs. “+”, or **Fig. 1C**: “Wt” vs. “KO”). To account for these technical variations, we included all necessary controls into each analysis session and abstained from directly comparing absolute lipid concentrations between different sessions.

### LC-MS/MS analysis

All samples were analysed using a Vanquish UHPLC system (Thermo Scientific #8362543) coupled to an ID-X Tribrid mass spectrometer (Thermo Fisher Scientific #FSN30160) equipped with a heated electrospray ionization (HESI) probe. For eicosanoid analysis, reversed-phase separation was performed on an Acquity UPLC BEH Shield RP18 column (2.1 × 100 mm, 1.7 µm; Waters) as previously described (PMID: 30050044). The mobile phase consisted of (A) Acetonitrile/water/acetic acid (60/40/0.02, v/v) and (B) Acetonitrile /IPA (50/50, v/v). The stepwise gradient conditions were carried out for 10 min as follows: 0–5.0 min, 1–55% of solvent B; 5.0–5.5 min, 55–99% of solvent B, and finally 5.5–6.0 min, 99% of solvent B. The flow rate was maintained at 0.5 mL/min, with an injection volume of 10 µL. Samples were kept at 4 °C during analysis, and the column temperature was set to 50 °C.Mass spectrometer parameters were as follow: ion transfer tube temperature, 300 ◦C; vaporizer temperature, 275 °C; Orbitrap resolution MS1, 120,000, MS2, 30,000; RF lens, 60%; maximum injection time MS1, 50 ms, MS2, Auto; AGC target MS1 and MS2 were Auto. Samples for analysis were run in both negative and 2500 V used for negative ion voltage; Aux gas, 10 units; sheath gas, 40 units; sweep gas, 1 unit. HCD fragmentation was performed using stepped collision energies of 15%, 25%, and 35%. Full-scan mode with data-dependent MS2 (ddMS2) was applied across a mass range of m/z 200–600. Internal calibration was performed using EASY-IC. A scheduled targeted inclusion mass list was used for the analysis, including 5-HETE, 5-KETE, 5-HETE-d8, and 5-KETE-d7. Peak identifications were performed using Excalibur 3.1 (Thermo Scientific, #OPTON-30965). The retention time, precursor ion mass/charge, and fragmentation patterns of 5-KETE, 5-HETE, 5-KETE-d7, and 5-HETE-d8 were confirmed using purchased standards. The following transitions were monitored: 5-KETE: 317.2122 > 203.1806, 5-HETE: 319.2280 > 115.0402, 5-KETE-d7: 324.2563 > 210.2242, 5-HETE-d8: 327.2783 > 116. 0463. For quantification, the area under the curve (AUC) for 5-HETE and 5-KETE was normalized to the AUC of the corresponding internal standards (5-HETE-d8 and 5-KETE-d7, respectively).

For the adult tail fin lipidomic analysis, 0.1 g of tail fin tissue from 3–6-month-old of *dhrs7^wt/wt^* and *dhrs7^mk218/mk218^* zebrafish were collected. The samples were sent to the UC San Diego Lipid Maps facility for LC-MS analysis. The tail fin tissues were homogenized in 1 mL 10% methanol in water suing a Bead Mill 24 (Fisher Scientific, 15-340-163). Internal standards were added to 100 μL homogenate. Eicosanoids were extracted using solid-phase extraction with Phenomenex Strata-X polymeric reversed-phase columns. The samples were dried and reconstituted in buffer A (water/acetonitrile/acetic acid, 60/40/0.02, v/v/v). Analysis was performed with a Water Acquity UPLC interfaced with an AB Sciex 6500 QTrap instrument. Chromatographic separation was achieved with a step gradient from 100% buffer A to 100% buffer B (acetonitrile/isopropanol, 50/50, v/v) over 5 minutes. Standard curves were generated under identical conditions. Data analysis was conducted using Analyst and MultiQuant software packages ^53^. The lipidomic data is presented in Table S1.

### Expression pattern of DHRS7 in different cell components

HEK293T cells from 10 plates were harvested and lysed as previously described. The nuclear pellet was collected by centrifugation at 1000× g for 10 min at 4℃. The supernatant was then transferred to a new tube, and the granule pellet was collected by centrifugation at 15,000 × g for 10 minutes at 4℃. The supernatant was transferred again to a new tube, and the microsome pellet was collected by high-speed centrifugation at 186,000× g for 1-2 hours at 4℃. The supernatant was used as the cytosol.

### NADPH production assay

For the NADPH production assay using different cell components, 10 μg of nuclear, granule, microsome and cytosol fraction were incubated with 1 μM 5(S)-HETE and 100 μM NADP^+^.

For the microsomal NADPH detection extraction experiments, *hs* DHRS7 (human), *dr* Dhrs7 (zebrafish), DHRS7B, DHRS7C, DHRS9, RDH11 were cloned and ligated to the pSBtet-eGFP vector. These constructs were transfected into HEK293K cells and selected using 10 μg/mL blasticidin S hydrochloride. Blasticidin-resistant cells were then treated with 1 μg/mL doxycycline for 24 hours, GFP-positive cells were sorted via flow cytometry to establish stable cell lines.

The enzyme assay was performed in a total volume of 100 μL containing reaction buffer (90 mM potassium phosphate, 40 mM KCl, 0.1 mM DTT, pH 7.5), the indicated amount of microsomes, 100 μM NADP^+^ or 100 μM NADPH, and 1 μM 5(S)-HETE or 1 μM 5-KETE. NADPH fluorescence was detected using Gen 5 plate reader with excitation at 340 nm and emission at 445 nm. The assay was conducted at 37℃ with readings taken every minute for 60-90 minutes.

### Determination of Michalis-Menten parameters

The first 20 timepoints of the NADPH kinetics were fitted with a line to approximate initial velocities (v_0_). The R^2^ values of each of these fits were averaged to yield an overall R^2^ value. Next the v_0_(NADP+) and v_0_(5(S)-HETE) isotherms were fitted with a Michaelis-Menten model v=V_max_*[S]/(K_m_+[S]) with [S] being the substrate concentration, V_max_ being the maximal enzyme velocity, and Km the Michaelis-Menten constant (indicating half-maximal enzyme saturation). Fitting and figure plotting was performed in MATLAB 2024b (Update 4,24.2.0.2833386, 64-bit, win64, MathWorks) using the base functions and the ‘Optimization’ and ‘Statistics and Machine Learning’ toolboxes. For better readability, the corresponding live script ‘MMcalculation.mlx’ was edited/annotated by ChatGPTo1 (OpenAI). This script obtained all numerical NADPH(t) data from a formatted Excel file (MichaelisMenten.xlsx), in which the NADPH concentration data was provided (in μM) in replicates as columns for each substrate concentration. For more details, please refer to the annotated MATLAB script.

### Spinning disk confocal imaging

For live imaging, 2.5-3 dpf anesthetized Tg(*lyz*:pm2-mKate2) and Tg(*mpeg1*: eGFP) larvae were amputated in Isotonic E3 tricaine, and immobilized in ∼200 μL of 1% low-melting agarose (Gold Biotechnology, A-204-100) dissolved in Isotonic E3, placed in glass bottom dishes (MatTek Corporation, P35G-1.5-20-C). After solidification of the agarose, the larvae were covered with 2-3 mL Isotonic E3 Tricaine to prevent and maintain anaesthesia during imaging. After 5 minutes of imaging, the buffer was replaced with the indicated medium for 1-hour imaging sessions. The medium used included Isotonic E3 tricaine with 7.5 μM Arachidonic acid (Sigma, A3611), 5 μM 5(S)-HETE (Cayman, 34230) or 1 μM 5-KETE (Cayman, 34250).

Imaging was performed at 28℃ in a heated imaging chamber (TOKAI HIT, WPI inc) using an inverted Nikon Eclipse Ti microscope. The microscope was equipped with a CFI Plan Apochromat Lambda D 10 × Objective lens (NA 0.45), a motorized stage, a Yokogawa CSU-W1 Spinning Disk unit, a Photometrics Prime BSI Scientific CMOS camera (2×2 binning), and NIS imaging software (Nikon, 5.11.01). Fluorescence emission was excited using 488 or 561 nm diode lasers (Nikon). Channel acquisition intensities/exposure times were set at 50%/100 ms for the 488 nm laser power or 80%/100 ms for the 561 nm laser power to detect either macrophages or neutrophils, respectively. Emission was collected using 525/36 (green) or 605/52 (red) bandpass filters with a high-sensitive sCMOS camera using band-pass dichroic mirrors (Chroma Technology Corp., 89100bs) for filtering two separate fluorescence emission spectra (525/36 and 605/52, Chroma Technology Corp., 89000 Sedat Quad) placed in front of the detector for isolated detection of green and red fluorescence, respectively.

### AlphaFold2 predictions of DHRS7 and hypothetical membrane positioning

AlphaFold2 structures from the listed species in S2A were superimposed in UCSF chimera (1.18) ^54^ by simultaneously fetching the structures and running Needleman-Wunsch alignment and homology matching algorithm with default settings ^55,56^. The predicted structure of hs DHRS7 was obtained from AlphaFold2 (ID:AF-Q9Y394-F1-v4), and the PDB file was used for membrane protein system and six steps was followed to assemble the hypothetical model using CHARMM36m and CHARMM-GUI. The protein structure was corrected and adjusted for lipid force fields and protonated with system pH 7.4 ^57^ the membrane positioning was determined using PPM 2.0. The system size for lipid bilayer was projected to heterogeneous lipid composition resembling that of ER membranes ^58^. The obtained model was then visually processed in UCSF Chimera (1.18).

### Statistics

No statistical methods were employed to determine the sample size, which aligns with those reported in previous publications ^22,23^. Statistical analyses were performed by using GraphPad Prism 8 (version 8.3.0) or MATLAB 2024b (Update 4,24.2.0.2833386, 64-bit, win64, MathWorks). Shapiro-Wilk test was used for measuring normal distribution for each dataset. Only if the data were normally distributed, a parametric unpaired two-tailed Welch’s t test was applied to the dataset. If the data were not normally distributed, non-parametric unpaired two tailed Mann-Whitney U test or ordinary one-way ANOVA (uncorrected Fisher’s LSD post-hoc test, two-tailed) were used. P values <0.05 were considered significant. All error bars represent the standard deviation (SD). Detailed sample sizes are listed in the figure legends. Animal experiments conducted on different days were aggregated. Experiments were not randomized, and investigators were not blinded to allocation during experiments or outcome assessment. For each treatment, both treated and control embryos were sourced from the same egg spawn. Animals were not reused across experiments. Larvae were selected based on normal morphology, a beating heart, and circulating red blood cells. All images in the figures represent typical phenotypes and expression patterns for their respective conditions. All detailed statistical tests are presented in Table S3.

## Acknowledgments

The research has been funded by the NIH grants and R35GM140883, and a BRIA award to P.N., a Tow fellowship to K.L.H., a Marie-Josée Kravis Women in Science Endeavor (Wise) fellowship to Y.N.M., and an experimental Immuno-oncology Scholars fellowship to M.L. Core facility services were in part funded by the NIH/NCI Cancer Center Support grant P30CA008748. We would like to thank Xuejun Jiang for valuable comments on the manuscript.

## Author contributions

Y.N.M., K.L.H., and P.N. conceived the project. K.L.H. and Y.N.M. conducted preliminary investigations to identify DHRS7 as the 5-HEDH candidate in A549 cells. K.L.H. generated the *dhrs7*^mk218/mk218^ zebrafish line. Y.N.M. generated the *dhrs7*^mk219/mk219^ zebrafish line, established DHRS7 HEK293T and A549 KO cell lines. Y.N.M. designed and performed all experiments in the Figures of this manuscript, except for LC-MS analysis, which was conducted by Y.A. from the T.B. and R.F. lab. M.L. provided *oxer1*^mk214/mk214^(*hcar1-4*) macrophage zebrafish. D.J.L. advised on ferroptosis treatments. Z.G. conducted structural simulations, advised on protein purification, assisted with manuscript editing, and helped preparing the figures. Y.N.M., and P.N. wrote the paper.

---

**Figure S1.**
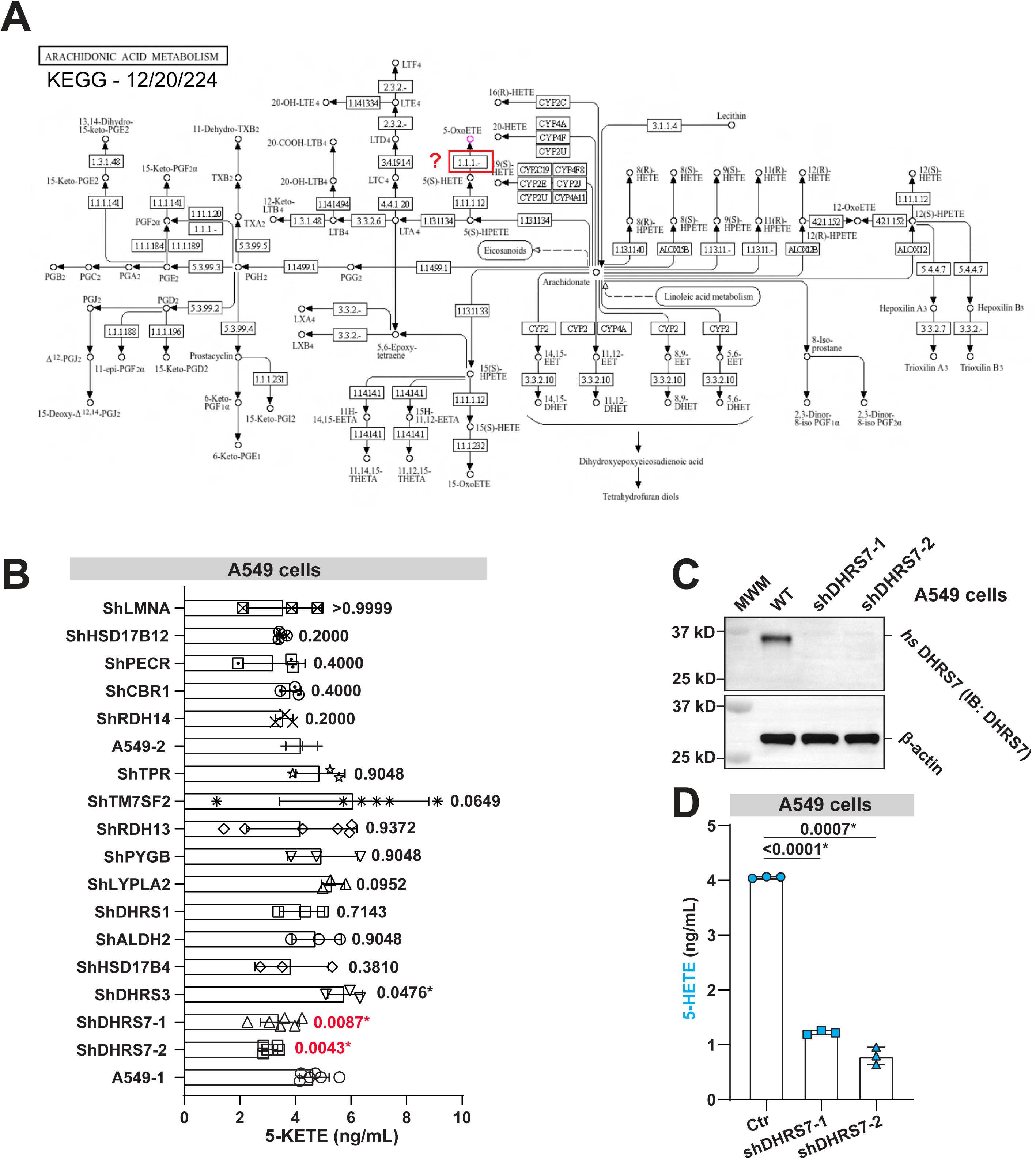
5-HEDH candidate screen. Related to Figure 1. **(A)** KEGG pathway scheme of arachidonic acid pathway showing the position of 5-HEDH (red). **(B)** Limited candidate screen for 5-HEDH using shRNA gene interference and LC-MS in A549 cells. Lipids extractions from shDHRS7-1, shDHRS7-2, shDHRS3, shHSD17B4, shALDH2, shDHRS1, shLYPLA2, shPYGB, shRDH13, shTM7SF2, shTPR were compared to A549-1, while shRDH14, shCBR1, shPECR, shHSD17B12, shLMNA were compared to A549-2. P values, unpaired, nonparametric, two tailed Mann-Whitney Test. **(C)** Western blot confirming DHRS7 knockdown by the two shRNAs (shDHRS7-1 and shDHRS7-2) used in the screen. β-actin was used as an internal control. MWM, molecular weight marker. **(D)** LC-MS assay showing 5-HETE production in DHRS7 knockdown A549 cells upon incubation with 1 μM 5-KETE. P values, unpaired, parametric, two tailed Welch’s t test. Asteriks, highlight significant changes as compared to respective control (p<0.05).

**Figure S2.**
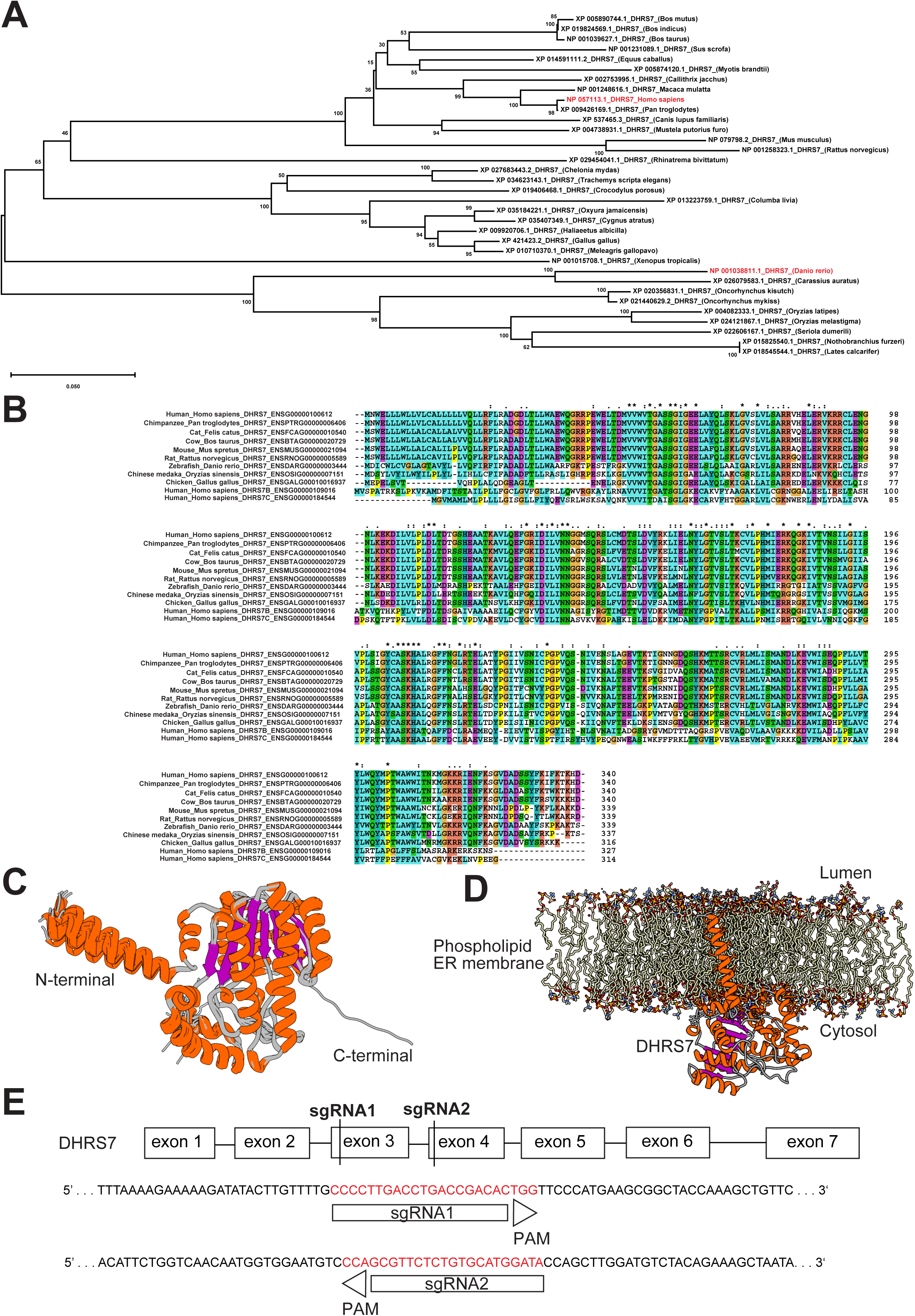
Phylogenetic analysis of DHRS7. Related to Figure 1. **(A)** Phylogenetic tree analysis of DHRS7 including human (*homo sapiens*), zebrafish (*danio rerio*) and 32 other species. **(B)** Protein sequence alignment of human DHRS7, DHRS7B, DHRS7C, zebrafish DHRS7 and orthologs from 7 other species. Asterisks (*), fully conserved residues. Colons (:) strongly similar conserved residues. Periods (.), weakly similar conserved residues. **(C)** Superimposed AlphaFold2 structures of 34 DHRS7 orthologs (shown in S2A). The colour key corresponds to alpha-helices (orange), beta-sheets (purple) and loops or unstructured (grey). **(D)** Cartoon model depicting the predicted structure of DHRS7 (ID:AF-Q9Y394-F1-v4), anchored in simulated ER phospholipid membrane leaflets **(E)** Schematic representation of DHRS7 CRISPR design. Two sgRNAs were designed to target exon 3 and exon 4.

**Figure S3.**
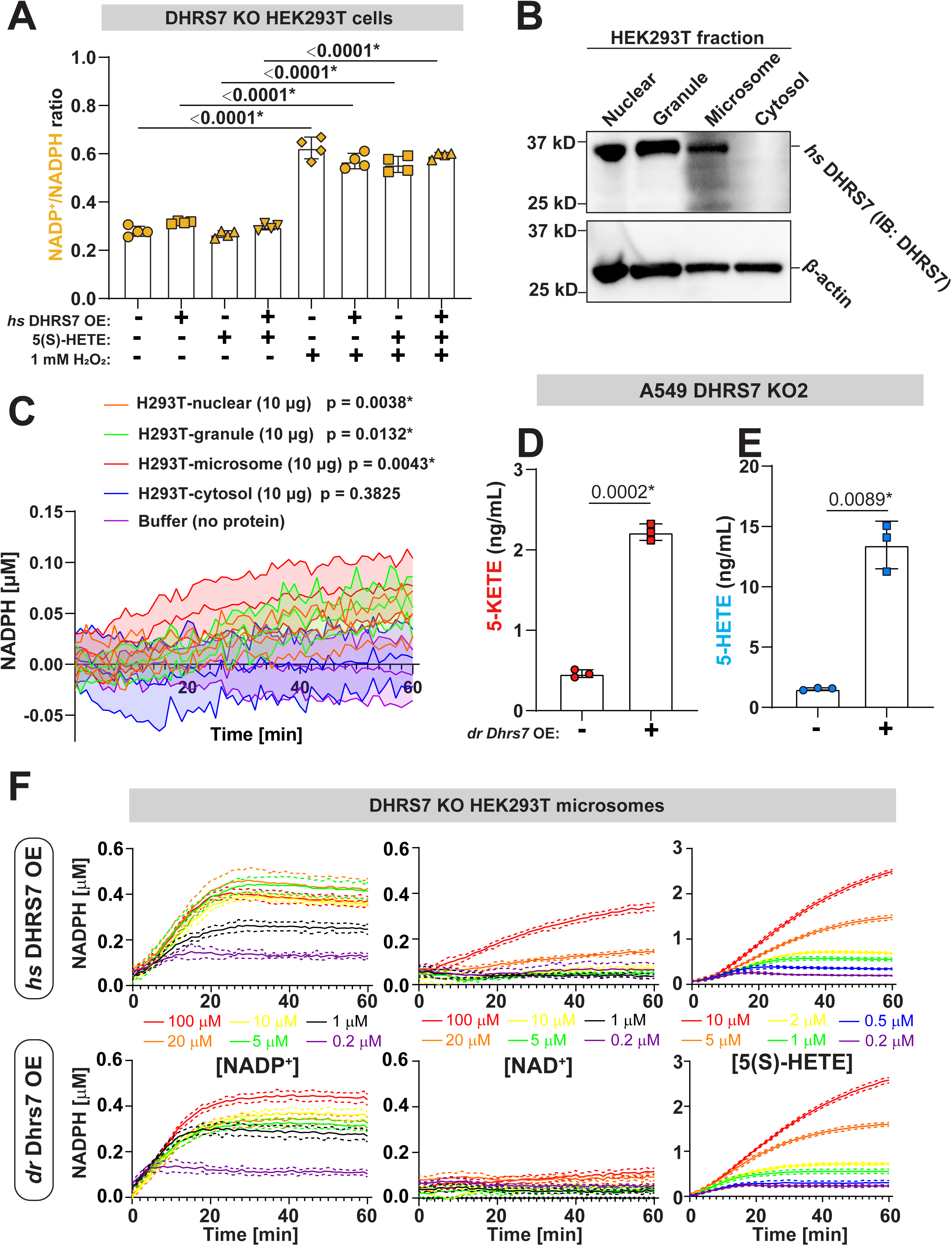
Extended characterization of DHRS7 as 5-HEDH, related to Figure 1-3. **(A)** NADP^+^/NADPH ratios in HEK293T DHRS7 KO cells ± *hs* DHRS7 OE, 1 μM 5(S)-HETE, or 1 mM H_2_O_2_ (20 min). P values, one-way ANOVA (uncorrected Fisher’s LSD test, two-tailed). **(B)** Western blot assay for endogenous DHRS7 protein expression in nuclear, granule, microsome, cytosolic fractions of HEK293T cells. β-actin was used as loading control. **(C)** NADPH production assays probing for 5-HEDH activity in nuclear, granule, microsome and cytosolic fractions incubated with 1 μM 5(S)-HETE and 100 μM NADP^+^. Statistical comparisons were performed at the 60 min time point. P values (microsomes), unpaired, nonparametric two tailed Mann-Whitney test. P values (other fractions), unpaired, parametric, two tailed, Welch’s t-test. Shaded plot regions, SD. **(D-E)** LC-MS assays of 5-KETE or 5-HETE production in DHRS7-KO A549 cells ± *dr* Dhrs7-OE and lipid substrate incubation as indicated. Dhrs7 overexpression was induced with 1 μg/mL doxycycline. P values, unpaired, parametric, two tailed Welch’s t test. Error bars, SD. **(F)** NADPH (or NADH) kinetics with 1.0975 pmol microsomal (top) *hs* DHRS7 or (bottom) *dr* Dhrs7 at the indicated, variable concentrations of (left) NADP^+^, (middle) NAD^+^, or (right) 5(S)-HETE, and (left-middle) 1 μM 5(S)-HETE, or (right) 100 μM NADP^+^, respectively. The v_0_ approximation of Figure 3H is based on the first 20 timepoints of the data in the left and right panels. Solid lines, mean values of data; dotted lines, SD. Asteriks, highlight significant changes as compared to respective control (p<0.05).

**Figure S4.**
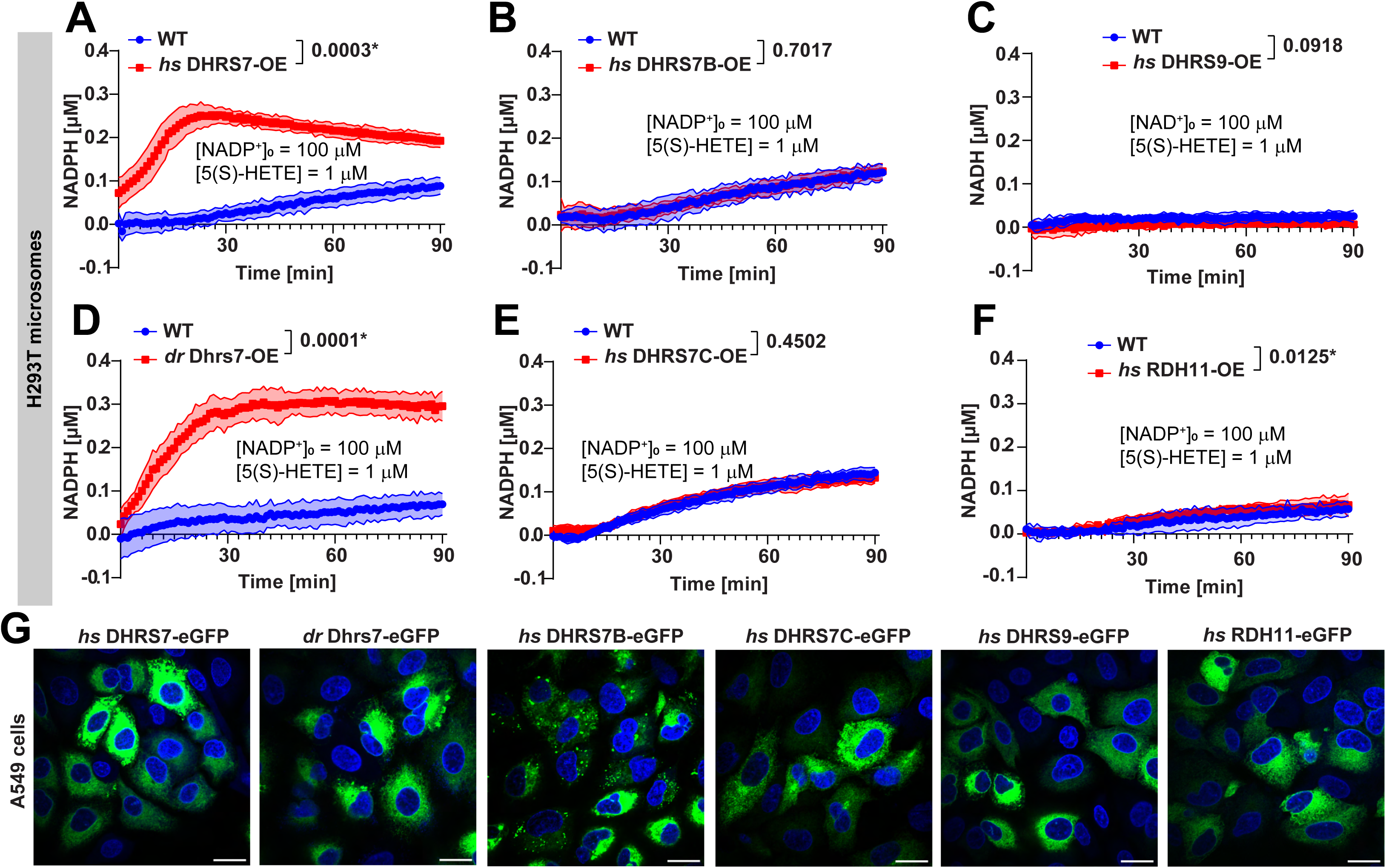
Extended characterization of DHRS7 as 5-HEDH, related to Figure 1-3. NADPH or NADH production assays with *wt* HEK293T microsomes overexpressing **(A)** *hs* DHRS7, **(B)** *hs* DHRS7B, **(C)** *hs* DHRS9, **(D)** *dr* Dhrs7, **(E)** *hs* DHRS7C, or **(F)** *hs* RDH11. 10 μg microsomes were incubated with 1 μM 5(S)-HETE and 100 μM NADP^+^, except for DHRS9 expressing microsomes, which were incubated with 1 μM 5(S)-HETE and 100 μM NAD^+^. NADPH or NADH levels are background corrected (using DMSO as vehicle control without substrate). P values, unpaired, parametric, two-tailed, Welch’s t test at the 20 min timepoint. Shaded plot regions, SD. **(G)** Representative images of the sub-cellular localization for the respective eGFP fusion proteins in A549 cells. Scale bar, 20 μm. Green, fusion protein. Blue, nucleus (DAPI). Asteriks, highlight significant changes as compared to respective control (p<0.05).

**Figure S5.**
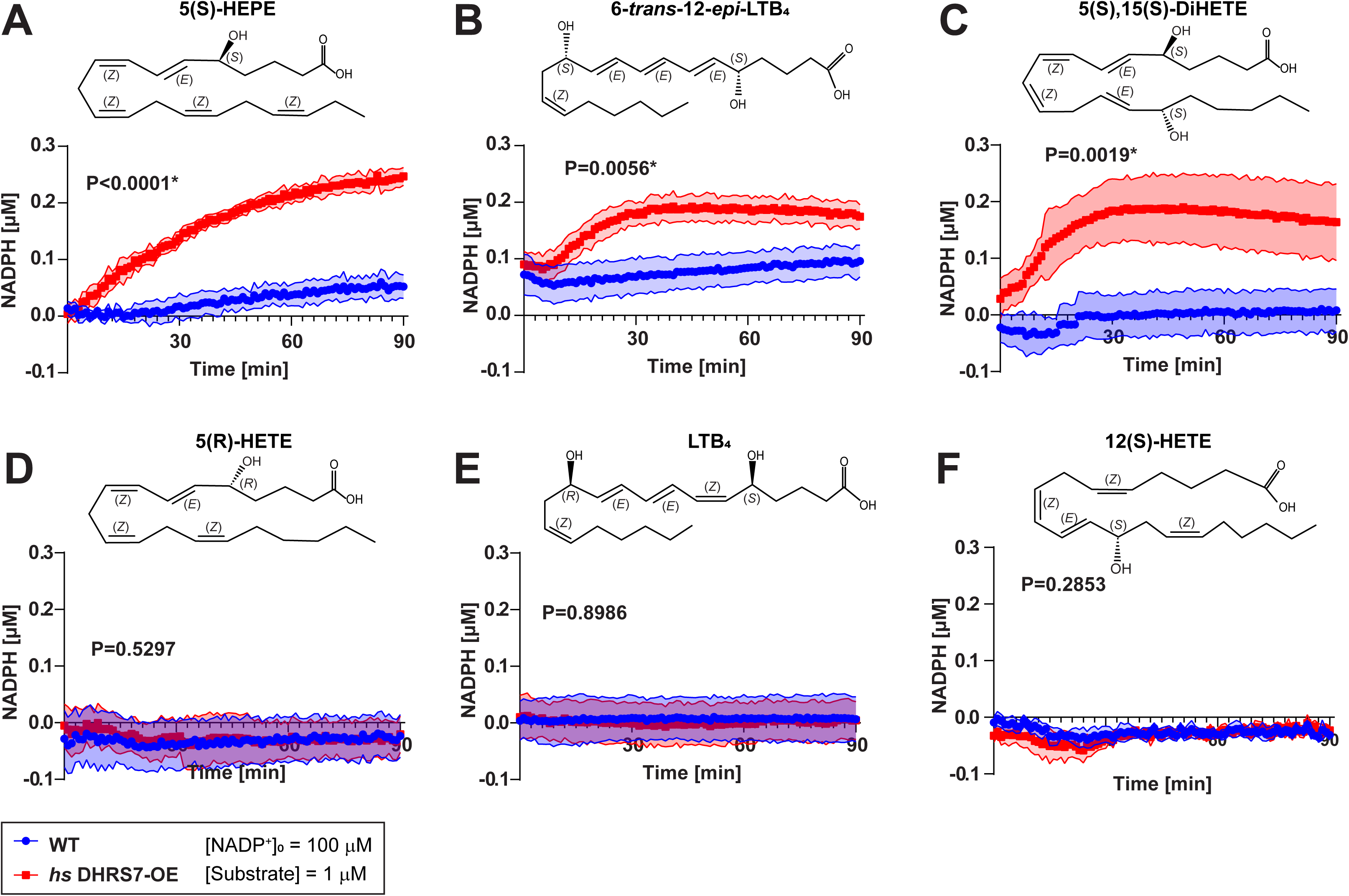
Extended characterization of DHRS7 as 5-HEDH, related to Figure 1-3. NADPH production assays were conducted with 100 μM NADP^+^ and 1 μM **(A)** 5(S)-HEPE, **(B)** 6-*trans*-12-*epi* LTB_4_, **(C)** 5(S),15(S)-DiHETE, **(D)** 5(R)-HETE, **(E)** LTB_4_, and **(F)** 12(S)-HETE. NADPH levels are background corrected (using DMSO as vehicle control without substrate). P values, unpaired, parametric, two tailed, Welch’s t test at the 20 min timepoint. Shaded plot regions, SD. Asterisks, highlight significant changes as compared to respective control (p<0.05).

**Figure S6.**
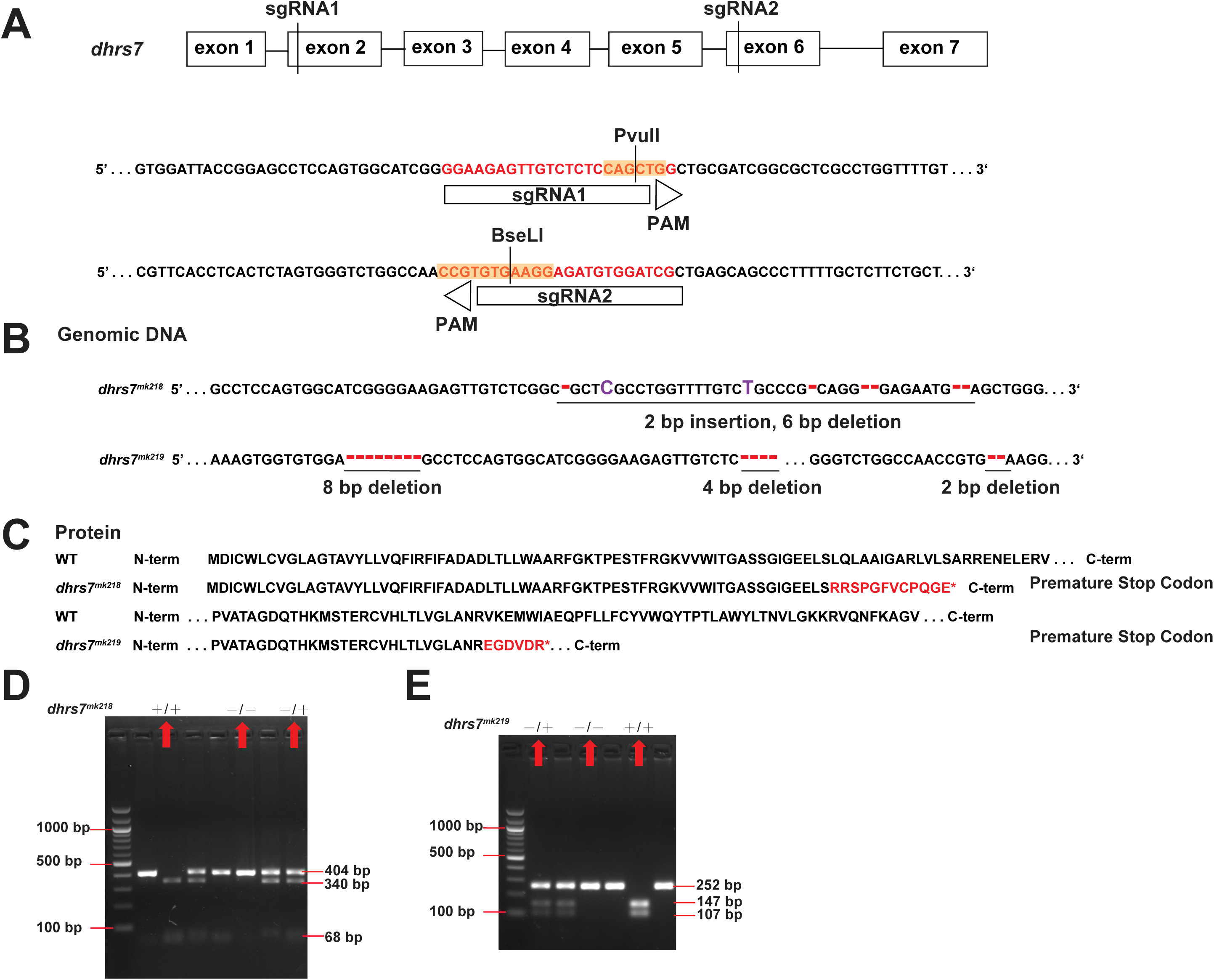
Generation of dhrs7 mutant zebrafish lines, related to Figure 4. **(A)** Schematic diagram showing the design of single guide RNAs (sgRNAs) to generate *dhrs7* CRISPR zebrafish. For *dhrs7^mk218/mk218^*, sgRNA1 was designed to target exon 2. Target disruption is predicted to occur at a PvuII restriction site (CAG^CTG) (highlighted by orange text) in the genomic DNA (deoxyribonucleic acid) sequence. For *dhrs7^mk219/mk219^*, two different sgRNAs were used together. Specifically, sgRNA2 was designed to target exon 6. Target disruption is predicted to occur at a BseLI restriction site (CCGTGTG^AAGG) (highlighted by orange text) in the genomic DNA sequence. **(B)** *dhrs7* mutant alleles. The mutant *dhrs7^mk218/mk218^* allele was generated by sgRNA1 and has a 4-base pair (bp) deletion resulting in a frameshift and the destruction of PvuII restriction site. The mutant *dhrs7^mk219/mk219^* allele was generated by co-injection of sgRNA1 and sgRNA2 and has a 12 bp deletion leading to the destruction of a PvuII restriction site, as well as a 2 bp deletion and the destruction of a BseLI restriction site, resulting in a frame shift. Purple text, mutant sequence (insertion) that differs from the *wt* sequence. Red text, deletion sequence that differs from the *wt* sequence. **(C)** The mutant *dhrs7^mk218^* and *dhrs7^mk219^* alleles both have premature stop codons in the coding sequence. Red text, mutant *dhrs7^mk218^* and *dhrs7^mk219^* sequences that differs from the *wt* sequence. **(D-E)** Heterozygous *dhrs7^mk218^* or *dhrs7^mk219^* zebrafish were crossed, and genomic DNA isolated from the progeny was PCR-amplified (PvuII-digested for *dhrs7^mk218/mk218^*, BseLI-digested for *dhrs7^mk219/mk219^*), and genotyped by agarose gel electrophoresis. As for *dhrs7^mk218/mk218^* (left panel), the PCR products from *dhrs7-1^wt^*^/*wt*^ fish are cleaved into two smaller products by PvuII (340 bp and 68 bp products). PCR products from homozygous *dhrs7-1*^*mk218*/mk218^ mutant fish are not cleaved by PvuII (404 bp product only). PCR products from heterozygote *dhrs7-1*^*wt*/mk218^ fish are identified by the presence of a cleaved *wt* allele and noncleaved mutant allele (404, 340 and 68 bp cleavage products). As for *dhrs7^mk219/mk219^* (right panel), the PCR products from *dhrs7-1^wt/wt^* fish are cleaved into two smaller products by BseLI (147 bp and 107 bp cleavage products). PCR products from homozygous mutant *dhrs7^mk219/mk219^* fish are not cleaved by BseLI (252 bp product only). PCR products from heterozygotes of *dhrs^wt/mk219^* were identified by the presence of a cleaved *wt* allele and non-cleaved mutant allele (252, 147 and 107 bp cleavage products).

**Figure S7.**
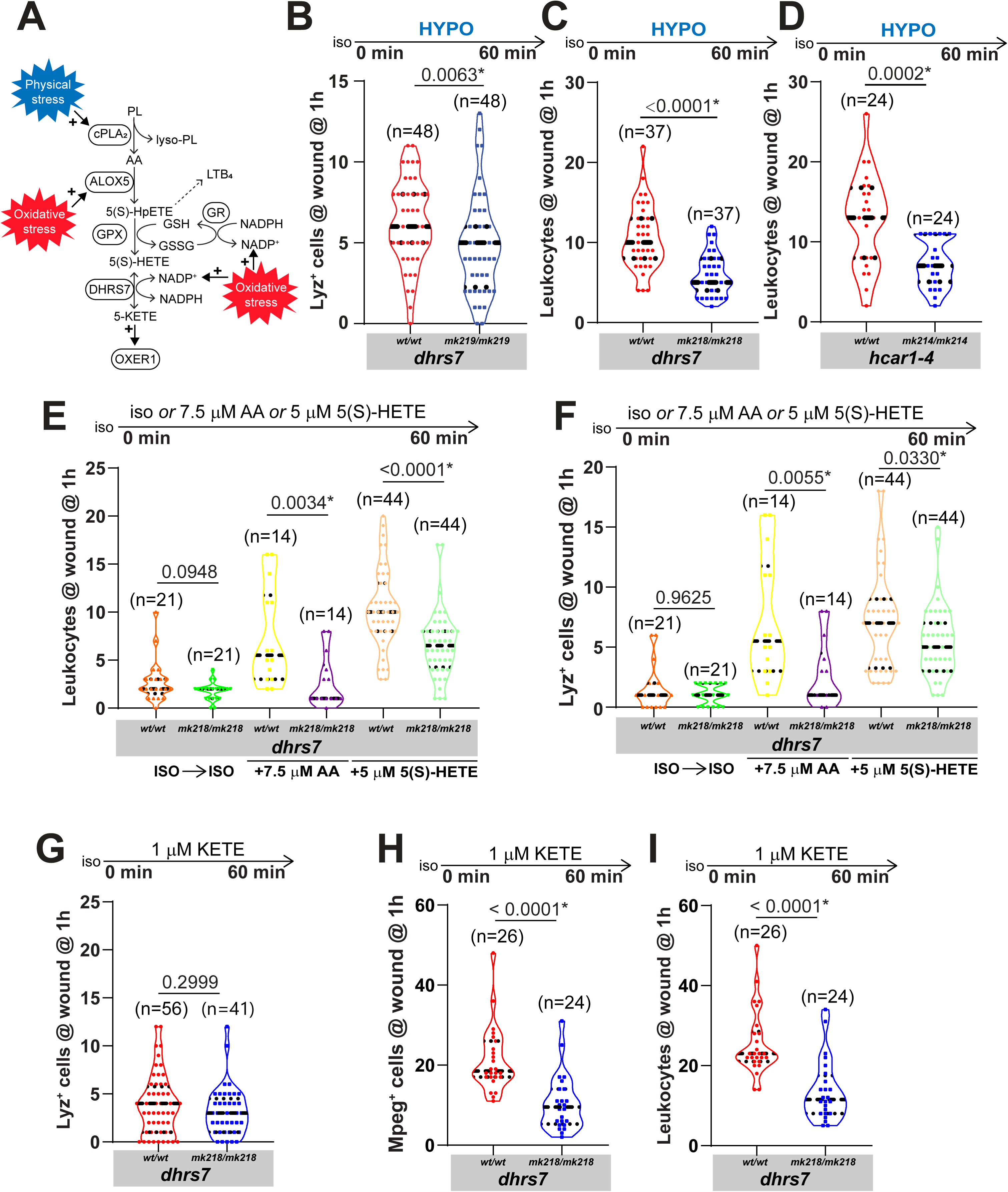
Characterization of Dhrs7 function in live zebrafish, related to Figure 4. **(A)** Schematic diagram depicting the oxoeicosanoid pathway. cPla_2_ is activated by physical stress (specifically, osmotic nuclear deformation) to release arachidonic acid (AA) from nuclear membrane phospholipids (PL). ALOX5 (5-lipoxygenase) metabolizes AA into 5-hydroperoxy-eicosatetraenoic acid (5(S)-HpETE). The catalytic site of ALOX5 is primed by free lipid peroxides (i.e., oxidative stress). 5(S)-HpETE may be either used for LTB_4_ production or reduced by a glutathione peroxidase (GPX) into 5-hydroxyeicosatetraenoic acid (5(S)-HETE). The respective oxidized glutathione (GSSG) is converted back to reduced glutathione (GSH) by glutathione reductase (GR) under consumption of NADPH and generation of NADP^+^. Under oxidative (=high NADP^+^) and physical stress (=high 5(S)-HETE), 5-KETE is generated from 5(S)-HETE. Under unstressed conditions, the reverse reaction is preferred. 5-KETE exerts its physiological effects through the OXER1 receptor (or its zebrafish ortholog Hcar1-4). **(B)** Quantification of Lyz^+^ neutrophils at the wound margins of *dhrs7^mk219/mk219^* or *wt* control siblings at 1 hour. **(C-D)** Quantification of “generic” leukocytes (identified by transmitted light contrast and motility) at the wound margins of **(C)** *dhrs7^mk218/mk218^* **(D)** *hcar1-4^mk214/mk214^* zebrafish compared to their respective control siblings (same experiment as in Figure 4C-D). **(E)** Quantification of Lyz^+^ neutrophils or **(F)** “generic” leukocytes at wound margin of *dhrs7^mk218/mk218^* mutant and sibling control larvae treated with the indicated lipids and isotonic bathing solution (Tg(*lyz*:pm2-mKate2) background). Quantification of **(G)** Lyz+ neutrophils, **(H)** Mpeg+ macrophages, or **(I)** “generic” leukocytes at the wound margins of *dhrs7^mk218/mk218^* mutant or *wt* sibling larvae after 1 hour of incubation with 1 μM 5-KETE in isotonic bathing solution (Tg(*lyz*:pm2-mk2) or Tg(*mpeg1*:eGFP) backgrounds). Dashed violin plot lines, first quartiles (top line), medians (middle line), third quartiles (bottom line). Parentheses, total number of wounded larvae. P values, unpaired, nonparametric, two-tailed Mann-Whitney U test. Note, (H-I) refer to the same experiment. Asterisks, highlight significant changes as compared to respective control (p<0.05).

## Movie Legends

**Video S1**. Live imaging of neutrophil recruitment to the wound margin of *dhrs7^wt/wt^* and *dhrs7^mk218/mk218^* larvae (tg(*lyz*:pm2-mk2) background), related to Figure 4C.

Representative time-lapse movie showing neutrophil recruitment to the wound margin. After 5-minute bathing in isotonic buffer, the bathing solution was switched to hypotonic buffer to trigger the wound response.

**Video S2**. Live imaging of neutrophil recruitment to the wound margin of *hcar1-4^wt/wt^* and *hcar1-4 ^mk214/mk21^*4 larvae (tg(*lyz*:pm2-mk2) background), related to Figure 4D.

Representative time-lapse movie showing neutrophil recruitment to the wound margin. After 5-minute bathing in isotonic buffer, the bathing solution was switched to hypotonic buffer to trigger the wound response.

**Video S3**. Live imaging of macrophage recruitment to wound margin in *dhrs7^wt/wt^* and *dhrs7^mk218/mk218^* larvae (tg(*mpeg1*:eGFP) background), related to Figure 4E.

Representative time-lapse movie showing macrophage recruitment to the wound margin. After 5-minute bathing in isotonic buffer, the bathing solution was switched to hypotonic buffer to trigger the wound response.

**Video S4**. Live imaging of macrophage recruitment to wound margin in *hcar1-4^wt/wt^* and *hcar1-4^mk214/mk214^* larvae (tg(*mpeg1*:eGFP) background), related to Figure 4F.

Representative time-lapse movie showing macrophage recruitment to the wound margin. After 5-minute bathing in isotonic buffer, the bathing solution was switched to hypotonic buffer to trigger the wound response.

## References

1. Forman, H. J. & Zhang, H. Targeting oxidative stress in disease: promise and limitations of antioxidant therapy. Nat. Rev. Drug Discov. 20, 689–709 (2021).

2. Bhattacharyya, A., Chattopadhyay, R., Mitra, S. & Crowe, S. E. Oxidative stress: an essential factor in the pathogenesis of gastrointestinal mucosal diseases. Physiol. Rev. 94, 329–354 (2014).

3. Pober, J. S., Min, W. & Bradley, J. R. Mechanisms of endothelial dysfunction, injury, and death. Annu. Rev. Pathol. 4, 71–95 (2009).

4. Niethammer, P., Grabher, C., Look, A. T. & Mitchison, T. J. A tissue-scale gradient of hydrogen peroxide mediates rapid wound detection in zebrafish. Nature 459, 996–999 (2009).

5. Yoo, S. K., Starnes, T. W., Deng, Q. & Huttenlocher, A. Lyn is a redox sensor that mediates leukocyte wound attraction in vivo. Nature 480, 109–112 (2011).

6. Schäfer, M. & Werner, S. Nrf2--A regulator of keratinocyte redox signaling. Free Radic. Biol. Med. 88, 243–252 (2015).

7. Holcombe, J. & Weavers, H. The role of preconditioning in the development of resilience: Mechanistic insights. Curr. Opin. Toxicol. 30, 100338 (2022).

8. Jelcic, M. & Niethammer, P. Do not scratch that mole! Trends Immunol. 36, 503–504 (2015).

9. Finkel, T. Signal transduction by reactive oxygen species. J. Cell Biol. 194, 7–15 (2011).

10. Sies, H. & Jones, D. P. Reactive oxygen species (ROS) as pleiotropic physiological signalling agents. Nat. Rev. Mol. Cell Biol. 21, 363–383 (2020).

11. Poole, L. B. The basics of thiols and cysteines in redox biology and chemistry. Free Radic. Biol. Med. 80, 148–157 (2015).

12. Tao, R. et al. Genetically encoded fluorescent sensors reveal dynamic regulation of NADPH metabolism. Nat. Methods 14, 720–728 (2017).

13. Molinari, P. E. et al. NERNST: a genetically-encoded ratiometric non-destructive sensing tool to estimate NADP(H) redox status in bacterial, plant and animal systems. Nat. Commun. 14, 3277 (2023).

14. Flores, M. V. et al. Dual oxidase in the intestinal epithelium of zebrafish larvae has antibacterial properties. Biochem. Biophys. Res. Commun. 400, 164–168 (2010).

15. Brothers, K. M. et al. NADPH oxidase-driven phagocyte recruitment controls Candida albicans filamentous growth and prevents mortality. PLoS Pathog. 9, e1003634 (2013).

16. Schoen, T. J. et al. Neutrophil phagocyte oxidase activity controls invasive fungal growth and inflammation in zebrafish. J. Cell Sci. 133, (2019).

17. Hogan, D. & Wheeler, R. T. The complex roles of NADPH oxidases in fungal infection. Cell. Microbiol. 16, 1156–1167 (2014).

18. De Deken, X., Corvilain, B., Dumont, J. E. & Miot, F. Roles of DUOX-mediated hydrogen peroxide in metabolism, host defense, and signaling. Antioxid. Redox Signal. 20, 2776–2793 (2014).

19. LeBert, D. C. & Huttenlocher, A. Inflammation and wound repair. Semin. Immunol. 26, 315–320 (2014).

20. Bennett, C. M. et al. Myelopoiesis in the zebrafish, Danio rerio. Blood 98, 643–651 (2001).

21. Redd, M. J., Cooper, L., Wood, W., Stramer, B. & Martin, P. Wound healing and inflammation: embryos reveal the way to perfect repair. Philos. Trans. R. Soc. Lond. B Biol. Sci. 359, 777–784 (2004).

22. Enyedi, B., Kala, S., Nikolich-Zugich, T. & Niethammer, P. Tissue damage detection by osmotic surveillance. Nat. Cell Biol. 15, 1123–1130 (2013).

23. Ma, Y., Hui, K. L., Gelashvili, Z. & Niethammer, P. Oxoeicosanoid signaling mediates early antimicrobial defense in zebrafish. Cell Rep. 42, 111974 (2023).

24. Powell, W. S. & Rokach, J. Targeting the OXE receptor as a potential novel therapy for asthma. Biochem. Pharmacol. 179, 113930 (2020).

25. Cossette, C. et al. Targeting the OXE receptor with a selective antagonist inhibits allergen-induced pulmonary inflammation in non-human primates. Br. J. Pharmacol. 179, 322–336 (2022).

26. Pochard, C. et al. PGI2 inhibits intestinal epithelial permeability and apoptosis to alleviate colitis. Cell. Mol. Gastroenterol. Hepatol. 12, 1037–1060 (2021).

27. Powell, W. S. & Rokach, J. The eosinophil chemoattractant 5-oxo-ETE and the OXE receptor. Prog. Lipid Res. 52, 651–665 (2013).

28. Kavanagh, K. L., Jörnvall, H., Persson, B. & Oppermann, U. Medium- and short-chain dehydrogenase/reductase gene and protein families : the SDR superfamily: functional and structural diversity within a family of metabolic and regulatory enzymes. Cell. Mol. Life Sci. 65, 3895–3906 (2008).

29. Geertz-Hansen, H. M., Blom, N., Feist, A. M., Brunak, S. & Petersen, T. N. Cofactory: sequence-based prediction of cofactor specificity of Rossmann folds. Proteins 82, 1819–1828 (2014).

30. Seibert, J. K. et al. A role for the dehydrogenase DHRS7 (SDR34C1) in prostate cancer. Cancer Med. 4, 1717–1729 (2015).

31. Uhlén, M. et al. A human protein atlas for normal and cancer tissues based on antibody proteomics. Mol. Cell. Proteomics 4, 1920–1932 (2005).

32. Stambergova, H., Skarydova, L., Dunford, J. E. & Wsol, V. Biochemical properties of human dehydrogenase/reductase (SDR family) member 7. Chem. Biol. Interact. 207, 52–57 (2014).

33. Araya, S. et al. DHRS7 (SDR34C1) - A new player in the regulation of androgen receptor function by inactivation of 5α-dihydrotestosterone? J. Steroid Biochem. Mol. Biol. 171, 288–295 (2017).

34. Rizzotto, A. et al. Reduction in Nuclear Size by DHRS7 in Prostate Cancer Cells and by Estradiol Propionate in DHRS7-Depleted Cells. Cells 13, (2023).

35. Jiang, M. et al. Characterization of the Zebrafish Cell Landscape at Single-Cell Resolution. Front. Cell Dev. Biol. 9, 743421 (2021).

36. Farnsworth, D. R., Saunders, L. M. & Miller, A. C. A single-cell transcriptome atlas for zebrafish development. Dev. Biol. 459, 100–108 (2020).

37. Zemanová, L., Kirubakaran, P., Pato, I. H., Štambergová, H. & Vondrášek, J. The identification of new substrates of human DHRS7 by molecular modeling and in vitro testing. Int. J. Biol. Macromol. 105, 171–182 (2017).

38. Minotti, G. & Aust, S. D. The requirement for iron (III) in the initiation of lipid peroxidation by iron (II) and hydrogen peroxide. J. Biol. Chem. 262, 1098–1104 (1987).

39. Katikaneni, A. et al. Lipid peroxidation regulates long-range wound detection through 5-lipoxygenase in zebrafish. Nat. Cell Biol. 22, 1049–1055 (2020).

40. Jiang, X., Stockwell, B. R. & Conrad, M. Ferroptosis: mechanisms, biology and role in disease. Nat. Rev. Mol. Cell Biol. 22, 266–282 (2021).

41. Erlemann, K.-R. et al. Regulation of 5-hydroxyeicosanoid dehydrogenase activity in monocytic cells. Biochem. J. 403, 157–165 (2007).

42. Patel, P. et al. Substrate selectivity of 5-hydroxyeicosanoid dehydrogenase and its inhibition by 5-hydroxy-Delta6-long-chain fatty acids. J. Pharmacol. Exp. Ther. 329, 335–341 (2009).

43. Powell, W. S., Gravelle, F. & Gravel, S. Metabolism of 5(S)-hydroxy-6,8,11,14-eicosatetraenoic acid and other 5(S)-hydroxyeicosanoids by a specific dehydrogenase in human polymorphonuclear leukocytes. J. Biol. Chem. 267, 19233–19241 (1992).

44. Enyedi, B., Jelcic, M. & Niethammer, P. The Cell Nucleus Serves as a Mechanotransducer of Tissue Damage-Induced Inflammation. Cell 165, 1160–1170 (2016).

45. Jelcic, M., Enyedi, B., Xavier, J. B. & Niethammer, P. Image-Based Measurement of H2O2 Reaction-Diffusion in Wounded Zebrafish Larvae. Biophys. J. 112, 2011–2018 (2017).

46. Shen, Z. et al. A synergy between mechanosensitive calcium- and membrane-binding mediates tension-sensing by C2-like domains. Proc Natl Acad Sci USA 119, (2022).

47. Venturini, V. et al. The nucleus measures shape changes for cellular proprioception to control dynamic cell behavior. Science 370, (2020).

48. Lomakin, A. J. et al. The nucleus acts as a ruler tailoring cell responses to spatial constraints. Science (2020).

49. White, R. M. et al. Transparent adult zebrafish as a tool for in vivo transplantation analysis. Cell Stem Cell 2, 183–189 (2008).

50. Nüsslein-Volhard, C. (Christiane) & Dahm, Ralf. Zebrafish: A Practical Approach (Practical Approach Series). 328 (Oxford University Press, 2002).

51. Miskolci, V. et al. Distinct inflammatory and wound healing responses to complex caudal fin injuries of larval zebrafish. eLife 8, (2019).

52. Ambaw, Y. A. et al. Profile of tear lipid mediator as a biomarker of inflammation for meibomian gland dysfunction and ocular surface diseases: Standard operating procedures. Ocul. Surf. 26, 318–327 (2022).

53. Quehenberger, O. et al. Lipidomics reveals a remarkable diversity of lipids in human plasma. J. Lipid Res. 51, 3299–3305 (2010).

54. Pettersen, E. F. et al. UCSF Chimera—a visualization system for exploratory research and analysis. J. Comput. Chem. 25, 1605–1612 (2004).

55. Jumper, J. et al. Highly accurate protein structure prediction with AlphaFold. Nature 596, 583–589 (2021).

56. Yang, Z., Zeng, X., Zhao, Y. & Chen, R. AlphaFold2 and its applications in the fields of biology and medicine. Signal Transduct. Target. Ther. 8, 115 (2023).

57. Klauda, J. B. et al. Update of the CHARMM all-atom additive force field for lipids: validation on six lipid types. J. Phys. Chem. B 114, 7830–7843 (2010).

58. Gee, S., Glover, K. J., Wittenberg, N. J. & Im, W. CHARMM-GUI Membrane Builder for Lipid Droplet Modeling and Simulation. ChemPlusChem 89, e202400013 (2024).

